# Fundamental transport mechanism of mucin-2 ER-to-Golgi trafficking identifies source of ER stress in inflammatory bowel disease

**DOI:** 10.1101/2024.05.13.593851

**Authors:** Margaretha A.J. Morsink, Lena S. Koch, Shixian Hu, Rinse K. Weersma, Harry van Goor, Arno R. Bourgonje, Kerensa Broersen

**Affiliations:** Department of Applied Stem Cell Technologies, TechMed Center, University of Twente, Enschede, The Netherlands; Department of Gastroenterology and Hepatology, University of Groningen, University Medical Center Groningen, Groningen, the Netherlands; Institute of Precision Medicine, The First Affiliated Hospital, Sun Yat-Sen University, Guangzhou, Guangdong, China; Department of Pathology and Medical Biology, University of Groningen, University Medical Center Groningen, Groningen, the Netherlands

**Keywords:** Mucin-2/ COPII-enlargement/ Inflammatory Bowel Disease/ ER-to-Golgi Trafficking/ TGF-β

## Abstract

The intestinal mucous layer relies on mucin-2 secretion. While the mucin-2 secretory pathway has been studied, endoplasmic reticulum (ER) to Golgi apparatus (Golgi) trafficking remains poorly understood. The size of mucin-2 exceeds the capacity of regular coat protein complex II (COPII) vesicles, responsible for ER-to-Golgi transport. After confirming conventional secretion of mucin-2, we showed that COPII vesicle enlargement is facilitated by TANGO1 and cTAGE5, and promoted by KLHL12. Inflammatory bowel disease (IBD) is characterized by a compromised mucous layer, altered activity of Transforming Growth Factor β (TGF-β), and increased ER stress. Using a cell culture, we showed that TGF-β inhibition induces TANGO1-mediated ER stress. Mucosal gene expression analysis in IBD patients confirmed elevated ER stress and validated concomitantly altered mRNA levels of TGF-β with mucin-2 and transport proteins TANGO1 and cTAGE5. In conclusion, we propose that the unsuccessful formation of enlarged COPII vesicles could be a source of ER stress in IBD, because of lowered TANGO1 protein expression, subsequently leading to decreased mucin-2 secretion.

## Introduction

The mucous membrane covers the entire epithelial lining of the human body facing the exterior, including the respiratory, digestive, and urinary tracts (1). In the intestines, mucin-2 is the predominant mucous protein, which is continuously secreted by intestinal goblet cells (2). Besides acting as the first line of defense against invading bacteria and microorganisms, the mucous layer provides a nutritional adherence site for commensal gut bacteria (3). Moreover, the mucous layer prevents dehydration, damage from gastric juices, and mechanical damage due to the peristaltic movement of the intestines (4, 5). Impairment of this layer has detrimental effects, as can be observed in patients suffering from inflammatory bowel disease (IBD) (6, 7), and has been linked to possible defects in post-transcriptional processing and secretion of mucin-2 (8). After post-translational dimerization and N-glycosylation within the endoplasmic reticulum (ER) (9–12), mucin-2 is extensively O-glycosylated in the Golgi apparatus. It is then oligomerized and subsequently packaged into goblet cell-residing secretory granules (13, 14). Following secretion, mucin-2 absorbs water to form a gel-like sheet with a protective function. The formation of secretory granules and post-Golgi transport mechanisms of mucin-2 has been well described (15–18). However, the ER-to-Golgi transport mechanism of mucin-2 remains largely unknown.

Transport of proteins out of the ER can be described by a conventional pathway (reviewed by (19)) which enables ER-to-Golgi transport, or through unconventional secretion pathways (reviewed by (20)) where proteins bypass the Golgi altogether. Conventional protein secretion requires a 60-80 nm large coat protein complex 2 (COPII)-coated vesicle for transport, which consists of inner COPII proteins SEC23 and SEC24, and outer COPII proteins SEC13 and SEC31 (reviewed by (21)). Brefeldin A (BFA), which inhibits COPII-mediated ER-to-Golgi transport (22), and Monensin, which inhibits trans-Golgi transport (23), are the most widely used compounds to inhibit conventional protein secretion (24). Early research has shown inhibited secretion of mucin-2 upon the addition of BFA and Monensin, indicating mucin-2 is conventionally secreted involving COPII vesicle transport from ER to Golgi (25, 26). However, mucin-2 exhibits a hydrodynamic radius of 176 nm upon exiting the ER, exceeding the dimensions of regular COPII vesicles (15, 16). Therefore, additional mechanisms must be involved to accommodate the large cargo in the vesicles.

Pro-collagen I (PC1), an essential extracellular matrix protein, is a 300-400 nm fibril upon exiting the ER. Its COPII-mediated ER-to-Golgi transport has been well examined by various researchers. Electron microscopy revealed the formation of enlarged COPII vesicles around the PC1 fibril (27) and various proteins facilitating this mechanism have been established. Cullin-3 ubiquitin ligase complex containing Kelch like protein 12 (Cul3-KLHL12) was identified as a key contributor for the enlarged COPII vesicle for PC1 trafficking (28). It mono-ubiquitylates outer COPII protein SEC31, which drives the formation of an enlarged COPII vesicle. In addition, overexpression of *KLHL12* induced accelerated secretion of PC1 (27, 29, 30). Moreover, transport and Golgi organization protein 1 (TANGO1) has been shown to enable transport of bulky cargo (31). TANGO1 interacts with cutaneous T-cell lymphoma-associated antigen 5 (cTAGE5) at ER exit sites (ERES) mediating the formation of enlarged COPII (32). The complementary role of TANGO1 and cTAGE5 has been described to facilitate transport of PC1, albeit the exact mechanisms are incompletely understood (27, 29, 33). Mucin variant MUC26B is trafficked through a TANGO1-mediated enlarged COPII structure in *Drosophila* (34). Therefore, we hypothesized a similar mechanism may enable the COPII-dependent transport of mucin-2. We further hypothesized that pathologies characterized by mucus layer deficiencies such as IBD, encompassing ulcerative colitis (UC) and Crohn’s disease (CD), are associated with differential regulation of ER-to-Golgi trafficking of mucin-2. This could in turn account for increased ER stress levels observed during IBD (35).

In this study, we reveal that mucin-2 ER-to-Golgi trafficking follows a process analogous to PC1, occurring through COPII vesicles and facilitated by TANGO1 and cTAGE5. Additionally, KLHL12 shows a promotive function in expanding COPII vesicles, aiding in the trafficking of the bulky mucin-2 from the ER to the Golgi apparatus. Furthermore, we identify a regulatory role of TGF-β for TANGO1. This discovery suggests that impaired COPII vesicle formation and deficient mucin-2 secretion may contribute to the elevated ER stress observed in IBD, a condition also characterized by altered levels of TGF-β. Analysis of mucosal gene expression of colonic tissue from IBD patients further confirmed high ER stress associated with altered expression levels of TGF-β1, mucin-2 and COPII transport proteins.

## Results

### Mucin-2 is Transported in Large COPII Vesicles Facilitated by TANGO1 and cTAGE5, and Promoted by KLHL12

To confirm previously published data suggesting COPII involvement in mucin-2 trafficking (25, 26), we studied post-Golgi mucin-2 levels in the intestinal goblet cell representing cell line HT29-MTX. Exposure to Brefeldin A (BFA), and Monensin both reduced post-Golgi mucin-2 levels, without affecting cell viability **(Supplementary Figure 1, Supplementary Figure 2 A-E)**, with BFA reducing post-Golgi mucin-2 levels in a dose-dependent manner. Additional observations, such as the partial localization of mucin-2 with the outer COPII coat protein SEC31A and the decrease in post-Golgi mucin-2 upon silencing of SEC31A **(Supplementary Figure 3, Supplementary Figure 2 F-H)**, further provided support for the involvement of COPII in ER-to-Golgi mucin-2 transport.

Nevertheless, it remains enigmatic how the large dimensions of mucin-2 (diameter ∼350 nm) are accommodated by the limited size of COPII vesicles (60-80 nm). Previously, large PC1 was shown to be trafficked to the Golgi in enlarged COPII vesicles facilitated by a set of identified proteins (27). We aimed to establish whether mucin-2 ER-to-Golgi trafficking involved a similar enlargement mechanism. After analyzing the size of the SEC31A punctae **(Supplementary Figure 2H**), we determined the degree of colocalization of SEC31A punctae larger than 300 nm with mucin-2. Approximately 80% (27 out of 34 punctae) colocalized suggesting trafficking of mucin-2 in large COPII vesicles analogous to PC1.

Previously, TANGO1, cTAGE5, and KLHL12 were shown to facilitate the enlargement of COPII vesicles (**Figure 1A**) (27, 28, 33) while silencing of these chaperones resulted in decreased secretion of PC1 (36). To investigate the involvement of these proteins for mucin-2 ER-to-Golgi trafficking, TANGO1, cTAGE5, and KLHL12 were targeted for silencing using siRNA (**Supplementary Figure 4**) and levels of post-Golgi mucin-2 and SEC31A protein expression were measured using Western Blot. A significant decrease in post-Golgi mucin-2 levels was observed following the decrease in expression of the COPII-enlarging proteins, indicating their involvement in mucin-2 trafficking (**Figure 1B-C**) matching previous reports for PC1 (36). Moreover, a decrease in the level of expressed SEC31A was observed by silencing TANGO1 and KLHL12, but not for cTAGE5 (**Figure 1D-E**), consistent with published data (32). Given the requirement of TANGO1 and KLHL12 for mucin-2 trafficking, we hypothesized that overexpression of TANGO1 and KLHL12, using a TANGO1-FLAG and KLHL12-FLAG plasmid (**Supplementary Figure 5**), would result in an increased level of post-Golgi mucin-2. While overexpression of KLHL12 indeed induced an increase in post-Golgi mucin-2 levels, interestingly, TANGO1 overexpression had no effect (**Figure 1F-G**). The combination of TANGO1 and KLHL12 overexpression contributed to an increased post-Golgi mucin-2 level, although not significantly (p=0.501). The involvement of TANGO1 and KLHL12 in mucin-2 trafficking was further verified using immunofluorescence. Partial colocalization of TANGO1 and mucin-2 was observed (**Figure 1H**). Overexpression of KLHL12-FLAG and its detection by means of a FLAG-antibody showed colocalization with mucin-2 as well (**Figure 1I**). These results were analogous to immunofluorescence results obtained for PC1 (27), suggesting trafficking in large COPII vesicles. Altogether, these results suggest that mucin-2 is trafficked in enlarged COPII vesicles facilitated by TANGO1 and cTAGE5, and promoted by KLHL12.

**Figure 1:**
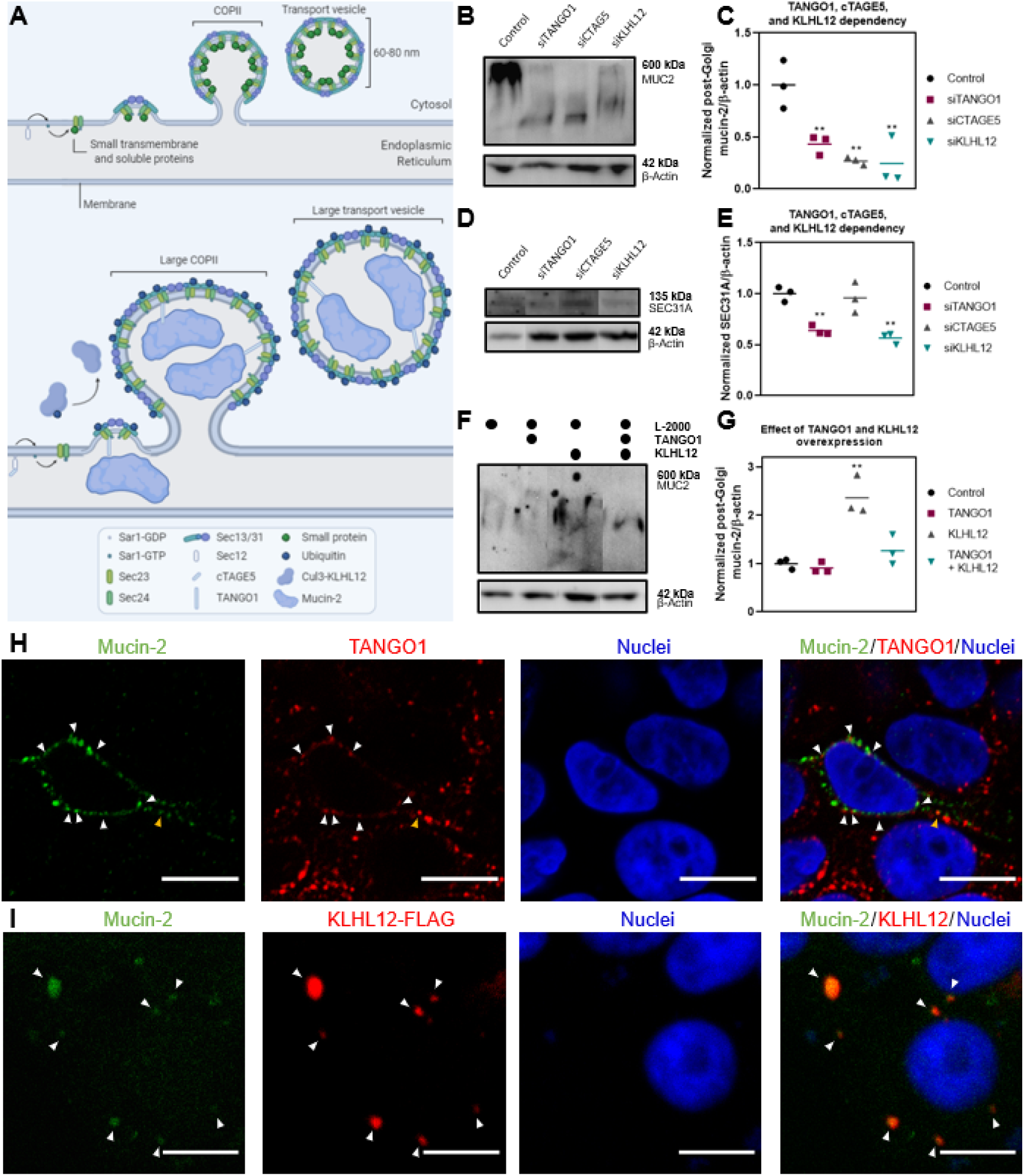
Mucin-2 transport in large COPII vesicles is promoted by KLHL12 and facilitated by TANGO1. A) A schematic representation of COPII formation for regular (top) and large (bottom) sized vesicles, showing the involvement of TANGO1, cTAGE5, and KLHL12. Created with BioRender.com. B) Western blot of post-Golgi mucin-2 after silencing of TANGO1, cTAGE5, and KLHL12. C) Semi-quantification of the Western Blot using ImageJ, comparing the signal intensity of mucin-2 to the signal intensity of β-actin, indicating a statistically significant decrease of post-Golgi mucin-2 levels upon silencing of TANGO1, cTAGE5, and KLHL12. D) Western blot of SEC31A after silencing of TANGO1, cTAGE5, and KLHL12. E) Semi-quantification of the Western Blot using ImageJ, comparing the signal intensity of SEC31A to the signal intensity of β-actin, indicating a statistically significant decrease of SEC31A expression upon silencing of TANGO1 and KLHL12, and similar expression after silencing cTAGE5. F) Western blot of mucin-2 upon overexpression of TANGO1 and KLHL12. G) Semi-quantification of the Western Blot using ImageJ, comparing the signal intensity of mucin-2 to the signal intensity of β-actin, indicating a statistically significant increase of post-Golgi mucin-2 levels upon overexpression of KLHL12 and a non-significant increase by overexpression of the combination of KLHL12 and TANGO1. H) Immunofluorescent staining of mucin-2 (green), TANGO1 (red), and nuclei (blue) showing partial colocalization of mucin-2 and TANGO1, as indicated by the white arrows. The yellow arrow indicates mucin-2 and TANGO1 colocalization further from the ER. Scale bar represents 10 µm. I) Immunofluorescence of mucin-2 (green), KLHL12-FLAG (red), and nuclei (blue), showing colocalization of mucin-2 and KLHL12-FLAG as indicated by the arrows. Scale bar represents 10 µm. Data was analyzed using one-way ANOVA followed by Bonferroni’s test. Significance was indicated as * (p < 0.05); ** (p < 0.01); *** (p <0.001) with respect to Control.

### TGF-*β* regulates TANGO1 protein expression

Recently, TGF-β was reported to upregulate TANGO1 protein expression to, in turn, facilitate PC1 secretion (36). As TANGO1 was found to facilitate trafficking to the Golgi for mucin-2, we predicted that exposure of HT29-MTX cells to TGF-β would mediate TANGO1-facilitated mucin-2 trafficking. We first investigated the effect of TGF-β1, as the most abundant isoform of the TGF-β family (37), on TANGO1 expression levels using Western Blot. In line with our expectations, TGF-β1 induced an upregulation of TANGO1 protein expression (**Figure 2B-C**) without affecting TANGO1 mRNA levels (**Figure 2N**), similar to results reported by Maiers and colleagues (36). Next, we sought if mucin-2 trafficking would be facilitated in response to TGF-β1-mediated TANGO1 upregulation. However, contradicting our expectations, we observed a significant decrease in mucin-2 post-Golgi levels (**Figure 2D-E**). *MUC2* transcription has been shown to converge with inflammatory pathways (38). The transcription factor NF-kB, a key regulator in inflammatory gene expression (39), is known to also play a positive role in *MUC2* transcription. IL-1β has been shown to induce NF-kB activity (40) while TGF-β1 is known as an inhibitor (7, 41). In light of this, it is possible that mucin-2 protein levels were inadvertently decreased as a function of TGF-β1 exposure (42). Using RT-qPCR we confirmed that *MUC2* mRNA levels were indeed severely downregulated by TGF-β1 (**Figure 2J**). We anticipated that these decreased *MUC2* mRNA levels upon exposure to TGF-β1 would be the result of TGF-β1-induced NF-kB downregulation. However, NF-κB levels were not altered by TGF-β1 (**Figure 3L**). Therefore, our attempts to rescue *MUC2* transcription using IL-1β (**Figure 2A, 2J**) were idle. Due to the low mRNA levels of *MUC2* following TGF-β1 stimulation, we anticipate reduced trafficking of mucin-2 from ER-to-Golgi, given there was less cargo to transport.

**Figure 2:**
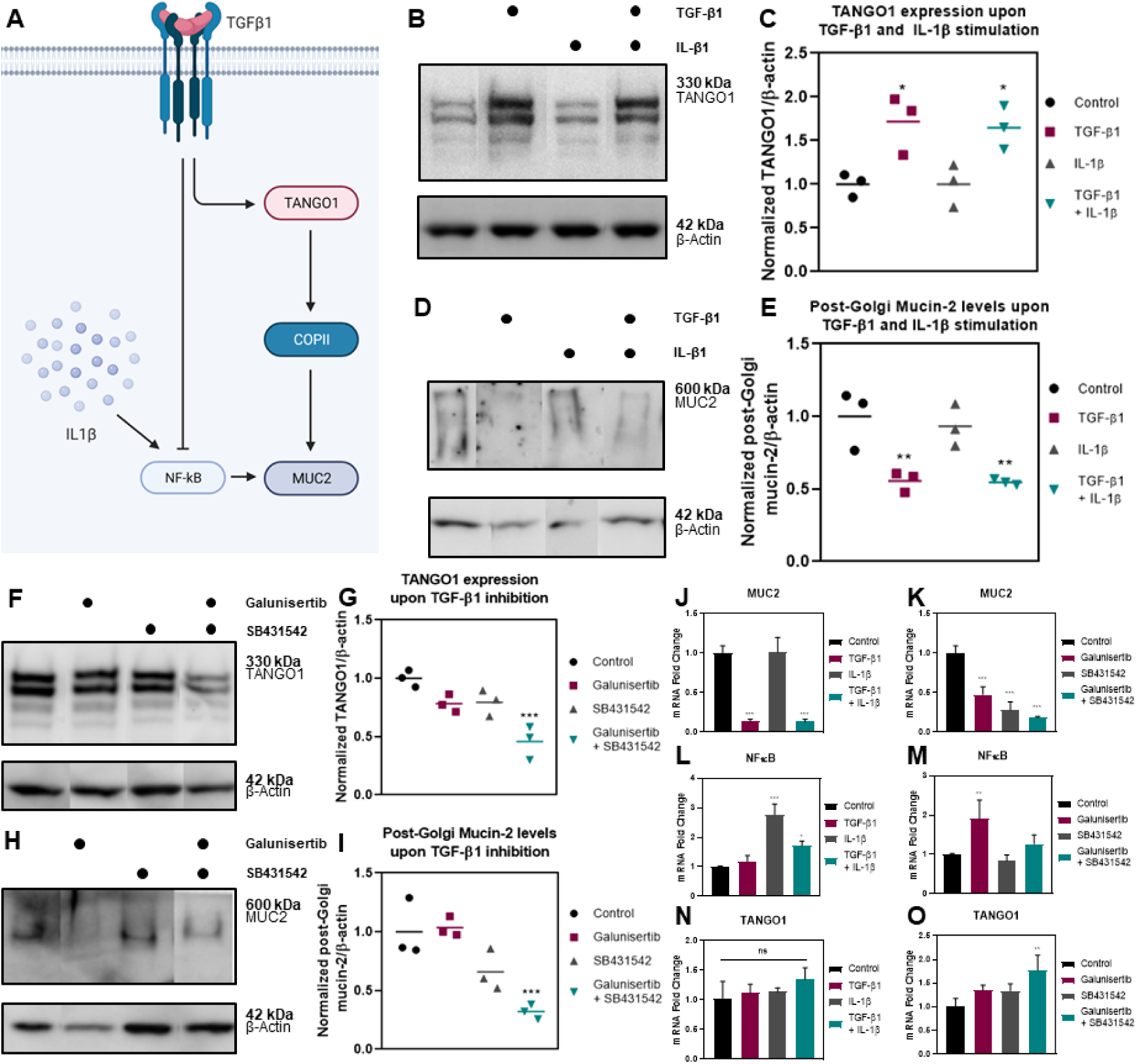
TGF-β facilitates mucin-2 trafficking through TANGO1. A) Schematic representation of the interaction of TGF-β1 with TANGO1, COPII, IL-1β, NF-κB, and MUC2. Created with BioRender.com. B) Western blot of TANGO1 after stimulation with TGF-β1 and IL-1β. C) Semi-quantification of the Western Blot using ImageJ, comparing the signal intensity of TANGO1 to the signal intensity of β-actin, indicating a statistically significant increase in TANGO1 expression upon addition of TGF-β1, and the combination of TGF-β1 and IL-1β. D) Western blot of post-Golgi mucin-2 after stimulation with TGF-β1 and IL-1β. E) Semi-quantification of the Western Blot using ImageJ, comparing the signal intensity of mucin-2 to the signal intensity of β-actin, indicating a statistically significant decrease of post-Golgi mucin-2 levels upon addition of TGF-β1, and the combination of TGF-β1 and IL-1β. F) Western blot of TANGO1 upon inhibiting TGF-β1 using Galunisertib and SB431542. G) Semi-quantification of the Western Blot using ImageJ, comparing the signal intensity of TANGO1 to the signal intensity of β-actin, indicating a statistically significant decrease of TANGO1 expression upon inhibition of TGF-β1 using Galunisertib and SB431542. H) Western blot of post-Golgi mucin-2 after inhibition of TGF-β1 using Galunisertib and SB431542. I) Semi-quantification of the Western Blot using ImageJ, comparing the signal intensity of mucin-2 to the signal intensity of β-actin, indicating a statistically significant decrease of post-Golgi mucin-2 levels upon inhibition of TGF-β1 using Galunisertib and SB431542. J-O) mRNA levels of MUC2, TANGO1, and NF-κB, normalized to GAPDH and control group using the 2^-ΔΔCt^ method determined with RT-qPCR. J) Decreased MUC2 mRNA levels upon addition of TGF-β1, and the combination of TGF-β1 and IL-1β. K) Decreased MUC2 mRNA levels upon inhibition of TGF-β1 using Galunisertib and SB431542. L) Increased NF-κB mRNA levels upon addition of IL-1β and TGF-β1 and IL-1β. M) Increased NF-κB mRNA levels upon inhibition of TGF-β1 through Galunisertib. N) TANGO1 mRNA levels after stimulation with TGF-β1 and IL-1β show no significant difference. O) TANGO1 mRNA levels upon inhibition of TGF-β1, showing significantly increased TANGO1 expression for the combination of Galunisertib and SB431542. Data was analyzed using one-way ANOVA followed by Bonferroni’s test. Significance was indicated as * (p < 0.05); ** (p < 0.01); *** (p <0.001) with respect to Control.

**Figure 3:**
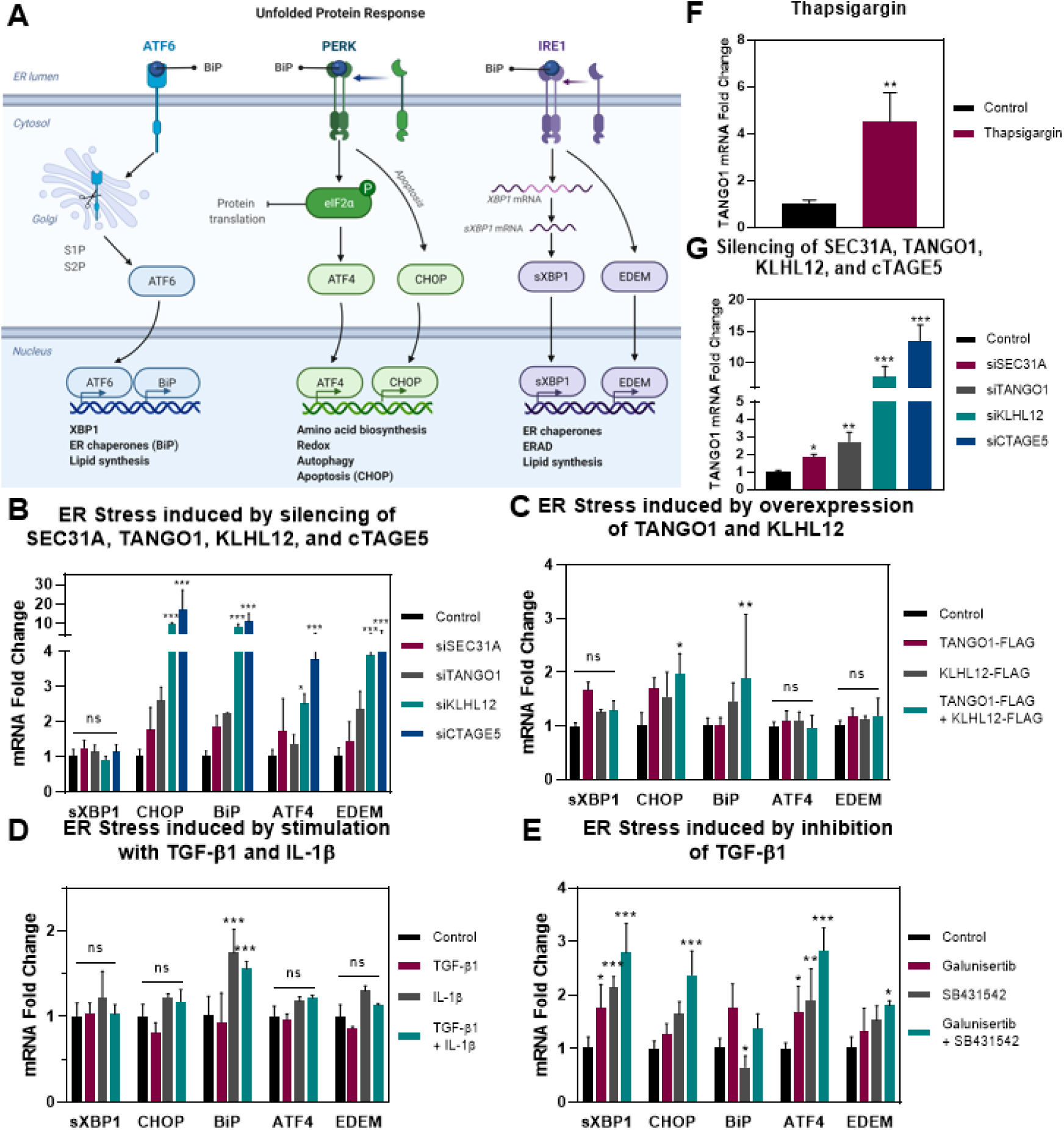
ER stress is increased by inhibiting the formation of large COPII vesicles. A) Schematic representation of the Unfolded Protein Response as measure of ER stress. The UPR consist of three pathways, namely ATF6, PERK, and IRE1. From each pathway, at least one target has been examined on mRNA level to indicate ER stress. Created with BioRender.com. B) ER stress is induced through silencing the formation of large COPII vesicles results in increased levels of ER stress in the CHOP, BiP, ATF4, and EDEM mRNA levels, but not by splicing of XBP1. C) ER stress is not induced by overexpression of TANGO1 and KLHL12 in sXBP1, ATF4, and EDEM. Elevated levels of CHOP and BiP are observed for the co-transfection of TANGO1 and KLHL12. D) ER stress as a result of exposure to TGF-β1 and IL-1β only shows elevated levels of BiP, yet other pathways are not altered as expected. E) ER stress is elevated upon inhibition of TGF-β1 by Galunisertib and SB431542. F) Thapsigargin upregulates TANGO1 mRNA levels. G) Silencing of SEC31A, TANGO1, KLHL12, and cTAGE5 upregulates TANGO1 mRNA levels. Data was analyzed using two-way ANOVA followed by Bonferroni’s test. Significance was indicated as * (p < 0.05); ** (p < 0.01); *** (p <0.001) with respect to Control of each specific gene tested.

As our experiments indicated that TGF-β1 plays a role in mucin-2 trafficking, we decided to inhibit TGF-β1 using the small molecules Galunisertib, SB431542, and their combination. Galunisertib is a small molecule inhibitor of the TGF-β1 receptor I, specifically downregulating the phosphorylation of Smad2, inhibiting activation of the canonical pathway of TGF-β (43). SB431542 also inhibits TGF-β1 receptor I, but, instead, by blocking phosphorylation of ALK4, ALK5, and ALK7 (activin receptor-like kinase) receptors (44, 45). In line with our expectations, TANGO1 protein expression was decreased significantly by inhibition of TGF-β1 (**Figure 2F-G**). In a response to the decreased TANGO1 protein levels, post-Golgi mucin-2 levels were also attenuated as measured by Western Blot (**Figure 2H-I**). This suggests the facilitating role of TGF-β1 in mucin-2 trafficking. However, we observed a decrease in *MUC2* mRNA levels as well (**Figure 2K**), unaffected by NF-κB mRNA levels (**Figure 2M**). Although *MUC2* mRNA levels were downregulated by Galunisertib, the post-Golgi mucin-2 levels were not, demonstrating low MUC2 mRNA levels still facilitate mucin-2 translation sufficient to study mucin-2 trafficking.

Due to the discrepancy between observed TANGO1 mRNA and protein levels in response to TGF-β1 stimulation, we investigated the TANGO1 mRNA levels in response to TGF-β1 inhibition (**Figure 2O**). We noted an upregulation of TANGO1 mRNA by inhibiting TGF-β1, directly contrasting the observed TANGO1 protein levels in response to TGF-β1 inhibition. The discrepancy between TANGO1 mRNA and protein levels suggests TANGO1 expression is mediated by post-transcriptional regulation (46, 47). Collectively, TGF-β1 plays an undeniable role in TANGO1 protein expression, which has been shown to facilitate mucin-2 secretion. However, due to the intricate role of TGF-β1 in various signaling pathways and cellular processes, we were not able to unambiguously report the effect on mucin-2 trafficking. The lack of TANGO1, as a result of TGF-β1 inhibition, severely impeded mucin-2 secretion, thereby suggesting the facilitating role of TGF-β1 in mucin-2 secretion.

### Inhibition of TGF-*β* leads to IBD-Associated ER Stress

Given the facilitating role of TGF-β1 in large COPII formation for mucin-2 trafficking, we hypothesized that the inhibition of TGF-β1 could be responsible for ER stress in goblet cells as has been observed in mucus-deficiency pathologies like IBD (35), due to compromised generation of large COPII vesicles. We examined several different genes in the Unfolded Protein Response (UPR) pathways through RT-qPCR (**Figure 3A**), previously described as a proxy for measuring ER stress (48). Thapsigargin stimulation served as a positive control (49). We showed elevated mRNA levels of spliced X-box Protein 1 (sXBP1), C/EBP-homologous protein (CHOP), immunoglobin binding protein (BiP), Activating Transcription Factor 4 (ATF4), and ER-degradation-enhancing-α-mannidose-like protein (EDEM) upon Thapsigargin stimulation (**Supplementary Figure 7**). To determine whether ER stress could be caused by the unsuccessful generation of large COPII vesicles, we silenced COPII protein SEC31A and COPII enlarging proteins TANGO1, KLHL12, and cTAGE5. Indeed, we observed increased ER stress levels upon inhibiting large COPII formation (**Figure 3B**), in line with published reports (50–52). Moreover, overexpression of TANGO1 and KLHL12, which facilitates trafficking in large COPII vesicles, did not elevate ER stress levels (**Figure 3C**). Collectively, these results elucidate the potential importance of the generation of large COPII vesicles to preclude ER stress.

Next, we investigated the effects of TGF-β1 and its inhibition on ER stress. Due to the positive effect of TGF-β1 on TANGO1 protein levels, therefore facilitating large COPII formation, we anticipated TGF-β1 to not induce ER stress. Indeed, no ER stress was observed upon stimulating the HT29-MTX cells with TGF-β1 (**Figure 3D**). Concomitantly, as the inhibition of TGF-β1 impedes large COPII formation, we predicted it would induce ER stress. In line with this expectation, elevated mRNA levels of all UPR targets were observed upon TGF-β1 inhibition (**Figure 3E**) consistent with previous reports (53, 54). These results demonstrate the importance of TGF-β1-mediated large COPII formation to inhibit generation of ER stress.

### TANGO1 is a Target of the Unfolded Protein Response

The observation that TANGO1 mRNA levels in response to TGF-β1 activation and inhibition (**Figure 2**) correlate with ER stress levels, prompted us to investigate TANGO1 mRNA levels of other ER stress situations. We hypothesized a UPR-mediated upregulation of TANGO1 mRNA levels as a result of ER stress. At the same time, we anticipated decreased TANGO1 protein levels through incomplete post-transcriptional regulation by low levels of TGF-β1. To address this hypothesis, we first confirmed increased TANGO1 transcription upon stimulation with the known ER stressor Thapsigargin in the goblet cell model HT29-MTX (**Figure 3F**). Moreover, silencing of COPII (enlarging) proteins SEC31A, KLHL12 and cTAGE5, which are known provokers of ER stress, showed a significant increase in TANGO1 mRNA levels as well (**Figure 3G**). Interestingly, silencing of TANGO1 somehow still upregulated TANGO1 mRNA levels (**Figure 3G**), but not protein levels (**Supplementary Figure 4**). Collectively, these findings suggest TANGO1 transcription is a target of the UPR. Further investigation of TANGO1 post-transcriptional regulation and clinical verification of TANGO1 mRNA and protein levels in IBD divulge new avenues for future research.

### Intestinal tissue from patients with IBD shows elevated ER stress levels and increased expression of MUC-2 trafficking genes

To validate our findings from the *in vitro* goblet cell model HT29-MTX, we investigated intestinal biopsy data from a cohort comprising individuals diagnosed with either UC or CD. We studied the expression profiles of genes involved in the UPR pathway, as well as components of the proposed ER-to-Golgi trafficking pathway in inflamed and non-inflamed regions of both the ileum and colon. In line with our *in vitro* findings, patients with UC and CD exhibited higher levels of ER stress compared to healthy controls with significantly increased mRNA levels of *XBP1*, *CHOP*, *BiP*, *ATF4* and *EDEM1* (**Figure 4A-E**). There was no difference in gene expression among the two disease states, however, ER stress was shown to be higher during active inflammation. We then evaluated the mucosal expression of mucin-2 and genes involved in its ER-to-Golgi trafficking. Compared to healthy controls, in patients with UC and CD, we observed a higher mRNA expression level of mucin-2 (**Figure 5A**). This was even more prominently increased in inflamed regions. Genes involved in mucin-2 ER-to-Golgi trafficking, such as SEC31a, TANGO1 and cTAGE5, were significantly increased in both UC and CD with a higher degree of expression during inflammation (**Figure 5B-D**). Against our expectations, *KLHL12* expression levels were not altered in both IBD subtypes compared to controls (**Figure 5E**). As a major negative regulatory element of mucosal inflammation, TGF-β1 levels (**Figure 5F**) were significantly increased in patients with IBD. The mucosal gene expression analysis of tissues from patients with IBD confirm that ER stress levels are increased in IBD and support our findings of differential expression of mucin-2 and its associated trafficking components.

**Figure 4:**
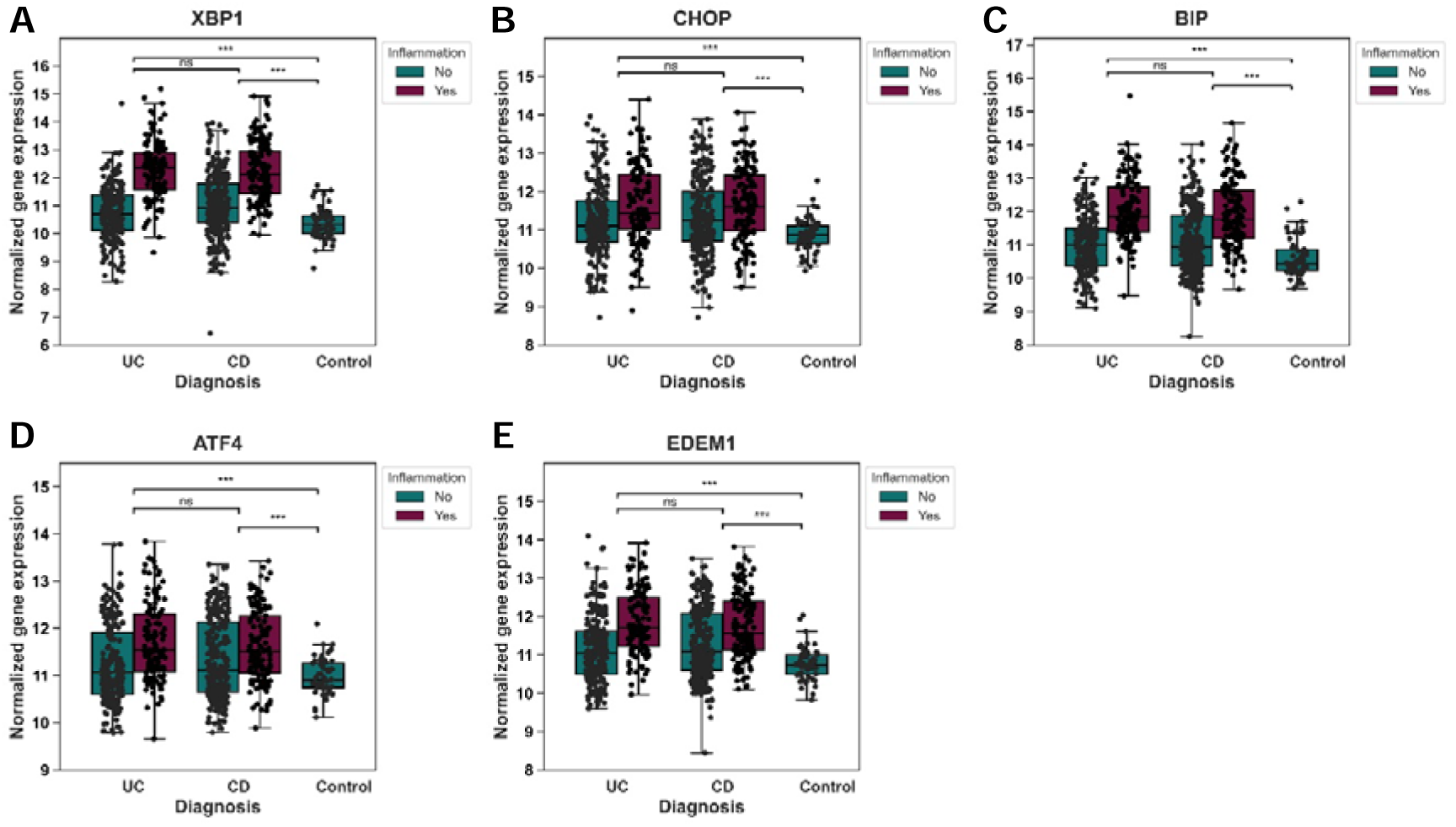
ER stress gene expression is increased in intestinal tissue from patients with ulcerative colitis (UC) and Crohn’s disease (CD). Intestinal biopsies from patients with IBD diagnosed with either UC or CD were sequenced and gene expression levels of ER stress markers XBP1, CHOP, BiP, ATF4 and EDEM1 (A-E) were determined. Gene expressions were compared between group of individuals with CD, UC and healthy controls using Mann-Whitney U-tests followed by post-hoc Bonferroni correction for multiple testing. Significance was indicated as * (p < 0.05); ** (p < 0.01); *** (p <0.001) with respect to healthy controls.

**Figure 5:**
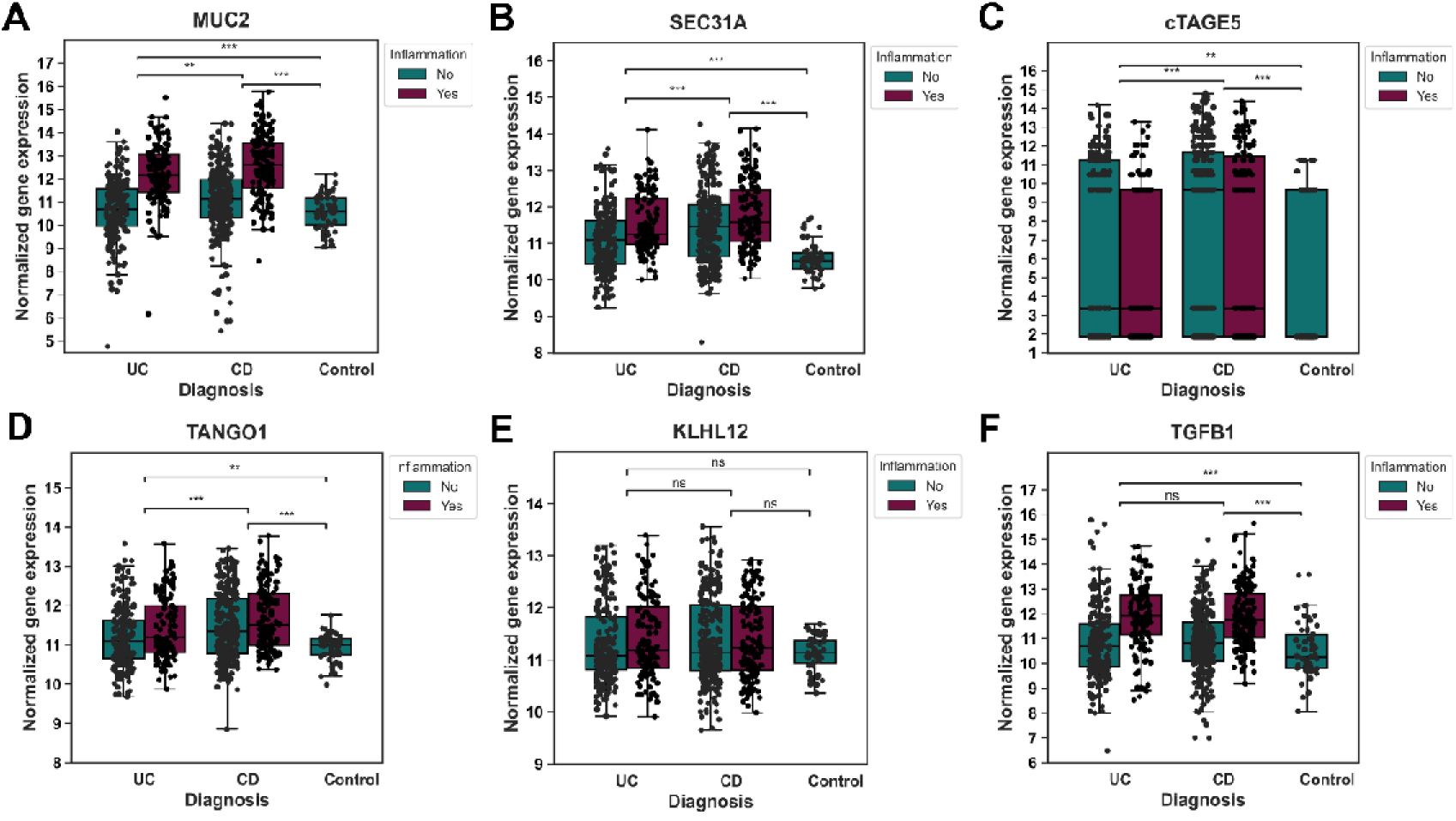
Gene expression data from patients with IBD possibly indicates an anti-inflammatory role of TGF-β and its regulatory effect on mucin-2 transcription and transport. Intestinal biopsies from an IBD patient cohort diagnosed with either UC or CD were sequenced and gene expression levels of mucin-2 and associated ER-to-Golgi transport factors in connection to TGF-β1 were evaluated. Patient data revealed elevated levels of MUC2 (A) along with genes involved in COPII-facilitated transport SEC31A (B), cTAGE5 (C) and TANGO1 (D). KLHL12 showed expression comparable to healthy controls. Upregulation of TGF-β1 (F) during UC and CD might divulge (reveal?) its role in promoting mucosal barrier function through amplified mucin-2 transcription and trafficking. Gene expressions were compared between group of individuals with CD, UC and healthy controls using Mann-Whitney U-tests followed by post-hoc Bonferroni correction for multiple testing. Significance was indicated as * (p < 0.05); ** (p < 0.01); *** (p <0.001) with respect to healthy controls.

## Discussion

This study reports on ER-to-Golgi trafficking of the mucin-2 protein, which is an important physiological component of the mucus layer lining the human intestines. Our results show that mucin-2 trafficking is COPII-dependent and is facilitated by proteins that have previously been documented to facilitate enlargement of COPII vesicles. We further show that TGF-β1 regulates TANGO1 protein expression and thereby facilitates COPII enlargement. We demonstrate that TGF-β1 inhibition is responsible for ER stress, in analogy to the ER stress as has been shown in IBD. Lastly, we validate our findings with IBD patient data which substantiate high ER stress levels and show high *MUC2* and associated COPII transport gene expression in association with upregulated TGF-β1 mRNA expression. Collectively, these findings indicate that …

### Mutations in COPII-Encoding Genes in Intestinal Diseases Illustrate COPII-Dependency of Mucin-2

Using BFA and Monensin, we demonstrated conventional trafficking of mucin-2 consistent with previous reports (25, 26, 55). Silencing of SEC31A further corroborated the COPII-dependency, which had not been shown for mucin-2 previously. Mutations in COPII genes have been linked to various pathophysiological defects in pro-collagen I (9, 56). However, no such direct links about COPII defects in mucin-2 related diseases are yet known, however, loss of TANGO1 protein levels is found in colon carcinomas, which are, in turn, associated with loss of mucous layer (47, 57). miRNA-429 has been indicated to downregulate inner COPII protein SEC23A and is associated with decreased mucin-2 secretion as well in a murine model of colitis (58), endorsing our findings of a COPII-mediated mucin-2 ER-to-Golgi transport mechanism. Interestingly, a clinical cohort study revealed single nucleotide polymorphisms (SNPs) in the SEC24B and SEC24B-antisense RNA 1 genes are associated with mucinous colorectal cancer (59) – a carcinoma which volume consists of at least 50% mucins (60). However, there is no evidence that proves that a defect in SEC24B affects mucin-2 secretion. Lastly, YIPF6 has been implicated in COPII vesicle formation, through interaction with inner COPII proteins SEC23 and SEC24 (61, 62), particularly facilitating the secretion of antimicrobial proteins in the intestines to maintain intestinal homeostasis. Mutations in YIPF6 cause spontaneous intestinal inflammation in mice and associated loss of mucin-2 was observed (63, 64). Although the exact mechanism through which YIPF6 induces colitis remains unknown, it is suggestive of mucin-2’s COPII dependency and COPII defects leading to pathophysiological disruptions in mucin-2 secretion.

Mutations in COP-II enlarging genes were furthermore implicated in various other diseases spanning cardiac (65, 66), neurological (67, 68) and liver (69, 70) defects, illustrating the consequences of disrupted ER-to-Golgi trafficking of various proteins. Therefore, future efforts should investigate the role and association of mutations in the COPII-enlarging proteins in intestinal diseases.

### Trafficking of Mucin-2 in Enlarged COPII Vesicles is Reminiscent of Transport of Other Large Proteins Including Pro-Collagen I

TANGO1 deficiency or knockout models have been associated with defects in secretory pathway genes and protein expression, as well as disrupted secretion of small and large cargo proteins including but not limited to PC1 (71). In vivo, TANGO1 knockdown in Drosophila decreased mucin-26B by disrupting COPII formation (34). TANGO1 and cTAGE5 have been implicated in transport of bulky chylomicrons in large COPII vesicles as well, and knockout of TANGO1 led to retained apoB within the cell (72). Moreover, KLHL12 was shown to colocalize with apoB and its silencing attenuated apoB secretion, corresponding to the results obtained for mucin-2 (73). Overexpressing TANGO and KLHL12 confirmed their promoting role for mucin-2 trafficking as has been observed for PC1 (28), collagen IV (74) and apoB (73) trafficking, through the mono-ubiquitylation of SEC31.

TANGO1’s facilitating role in formation of large COPII vesicles is unequivocal (31, 75), yet there are reports of its ambiguous mechanism in the generation of large COPII vesicles. Schekman and colleagues suggest a co-packaging of TANGO1 with the cargo (29), whereas Malhotra and colleagues demonstrated that TANGO1 does not leave the ER, but forms a ring around the COPII vesicle (33). The distanced colocalized punctae of TANGO1 and mucin-2 (**Figure 1B**, yellow arrow) suggest transport of TANGO1 away from the ER, which has been validated in recent results that show TANGO1 colocalizing with Sec16A indicating an ERES localization (71). In addition, TANGO1 might play another role in the mucin-2 secretory pathway occurring in the Golgi apparatus, as it has been shown to aid in O-glycosylation for MUC26B in *Drosophila* (34).

### TGF-*β* Facilitates Mucin-2 Trafficking Through TANGO1 Protein Expression, However, Molecular Mechanism of TGF-*β* on MUC2 Transcription Remains Enigmatic

TGF-β has been implicated in IBD as lower active levels are observed (76), due to increased expression of TGF-β inhibitor Smad7 in the intestinal mucosa (77, 78), prompting us to investigate the effect of TGF-β on mucin-2 trafficking. Consistent with previous research (36, 79), TGF-β induced expression of TANGO1. The effect of inhibiting TGF-β on TANGO1 has not been reported previously, however, the decreased expression is in line as an antagonistic response to TGF-β stimulation. Decreased post-Golgi mucin-2 levels resulting from TGF-β inhibition also endorse a facilitating role of TGF-β in TANGO1 and mucin-2 regulation. The specific role of TGF-β in mucin regulation remains ambivalent, however. In line with our findings on MUC2 mRNA levels, TGF-β was found to induce MUC5AC mRNA expression (80). At the same time, however, TGF-β was implicated in downregulation of MUC5AC expression (81). Moreover, a functional cooperation of TGF-β upregulation and NF-κB signaling was found to elevate MUC2 mRNA levels in response to the nontypeable Haemophilus influenzae bacteria (82). Others suggest an inhibitory effect of TGF-β to NF-κB (7, 41), yet our findings show no effect of TGF-β on NF-κB. IL-1β has been shown to activate transcription factor NF-κB (40), corresponding to our results. However, MUC2 transcription did not seem altered by NF-κB, contradicting numerous findings on MUC2 transcription (42, 83–88). Mucin-2 protein levels do not always correlate to MUC2 mRNA levels due to extensive post-translational modifications (89), as observed by the aberration in response to Galunisertib. Ultimately, the molecular mechanism through which MUC2 mRNA levels and mucin-2 trafficking are decreased in response to TGF-β exposure remains unknown.

The discrepancy between TANGO1 mRNA and protein levels suggests post-transcriptional regulation. Based on our findings related to TGF-β and its inhibition, we suspect it plays a role in TANGO1 post-transcriptional regulation, however we lack the evidence to support this statement. In general, the transcription and translation of TANGO1 have been poorly investigated (90). We have shown TANGO1 transcription is a target of the UPR, consistent with what is known about TANGO1 and other COPII proteins (36, 91). Other post-transcriptional and post-translational modifications of TANGO1 that have been reported including O-glycosylation (47) and phosphorylation (46), both impacting COPII formation. Various other phosphorylation sites of TANGO1 have been described, yet their exact mechanisms have not been investigated (50). Generally, post-transcriptional and post-translational modifications play an important role in the formation of large COPII vesicles (92). Recently, it was shown that TANGO1 expression was dependent on inflammasome activation of caspase-1, as well as signaling from IL-1 and TGF-β (79). Elucidating the post-transcriptional mechanism of TANGO1, possibly mediated by TGF-β, could identify therapeutic targets for IBD and fibrosis.

### TGF-*β* Inhibition Generates ER Stress in Intestinal Diseases Due to Unsuccessful Formation of Large COPII Vesicles

In addition to investigating the effects of TGF-β on mucin-2 trafficking, we examined its effect on other hallmarks of IBD pathogenesis and pathology, namely increased levels of ER stress in the goblet cells (93). There are several factors that could induce ER stress in IBD, although the exact mechanisms are not fully understood (94). Environmental factors, such as the interactions with the microbiota, are thought to contribute to ER stress (35). Moreover, mutations in UPR genes, such as XBP1, have been implicated in early onset Crohn’s disease (95), and mutations in the MUC2 gene, leading to unsuccessful O-glycosylation in the ER, have been shown to cause IBD-related ER stress (93, 96). Increased transcription of MUC2 has been shown to induce ER stress as well, due to the complex folding and post-translation processing of mucin-2 in the ER (97). Inflammatory states in the intestines induce mucin transcription and translation, thereby exerting a sudden increase in workload of the ER, leading to ER stress (98). Therefore, proper UPR signaling is essential to maintain ER homeostasis and cell functionality, as it attempts to restore balance. However, if protein accumulation at the ER lumen is beyond restoration, CHOP signaling is activated and induces apoptosis (48). Alterations in COPII genes have been shown to increase ER stress and activate UPR pathways in many cases (50–52), endorsing our findings. TANGO1 and KLHL12 overexpression were not shown to induce ER stress, elucidating their importance for ER homeostasis. TGF-β, and the subsequent increased TANGO1 protein expression, indeed did not affect ER stress levels, in contrast to findings of Maiers and colleagues (36), however in support of our theory that TGF-β enables mucin-2 transport. Concomitantly, TGF-β inhibition, and the decreased TANGO1 protein levels, did activate the UPR signaling pathways, supporting our theory.

Impaired TGF-β signaling is also found in Celiac Disease, also known as gluten intolerance, associated with elevated ER stress levels in the intestines and an altered mucosal layer (99–101), further suggesting the importance of TGF-β in maintaining ER homeostasis. Lastly, increased availability of IL-1β has been implicated in ER stress due to aberrant mucin formation, explaining our observed increased BiP levels (102). Collectively, we predict the increased Smad7 signaling inhibiting TGF-β in IBD contributes to the increased ER stress, due to a lack of TANGO1 protein availability, ultimately leading to decreased mucin-2 secretion.

### Patient Data Provide a Complex View on Regulation of Mucin-2 and Associated ER Stress

IBD is a complex and multifactorial disease caused by genetic, environmental, and microbial imbalances (35). The two major forms, ulcerative colitis (UC) and Crohn’s disease (CD) are both marked by a loss of mucus layer functionality (103). Corroborating our in vitro findings using the goblet cell line HT29-mtx, the mucosal gene expression analysis of intestinal tissue from patients with UC and CD showed significantly elevated ER stress levels in intestinal biopsies. Several studies have reported on increased ER stress in UC and CD intestinal tissues in connection with impaired mucin formation (102, 104, 105). Our patient data revealed significantly increased mucin-2 mRNA and its associated ER-to-Golgi trafficking genes. Mucin-2 mRNA and protein levels during IBD have been reported to vary. This was attributed to different origins of the disease, be it genetic or environmental, analysis methods and even patient cohort demographics (as reviewed in (106)). However, multiple studies agreed on an increase in MUC2 mRNA in UC and CD which was exacerbated in inflamed regions matching our findings (42, 102, 107–109). Furthermore, this human data confirms our earlier conclusions that increased expression levels of mucin-2 are concomitant with increased levels of SEC31A, cTAGE5 and TANGO1, signifying their association. In opposition to our discoveries, TGF-β1 mRNA expression levels were significantly elevated in UC and CD intestinal samples, now showing a positive relationship between TGF-β and TANGO1 mRNA levels paralleling the TANGO1 protein regulation by TGF-β we had established earlier. Naturally, patient data present a more holistic view on the molecular processes during IBD compared to our data derived from the in vitro goblet cell model HT29-mtx, also taking into account for complex immune system interactions during chronic and acute inflammation. While we resolved the pathway of ER-to-Golgi mucin-2 trafficking in an isolated goblet cell model during health and intestinal inflammation, the in vivo patient data shows a differential gene expression. We mimicked short term inflammation to reveal the role of TGF-β mediated mucin trafficking. The lack of an integrated immune component denotes one limitation of the in vitro study. Integrating LPMCs, PBMCs or dendritic cells could help elucidate the role of TGF-β mediated TANGO1 and subsequent mucin-2 regulation as they have been shown to be crucial for TGF-β signaling in the intestinal mucosa in inflammatory states (110–112).

Similar to mucin-2, TGF-β expression has been shown to vary in patients with IBD with a discrepancy of mRNA and protein levels (as reviewed in (113, 114)). The increased mRNA levels could be an indicator of its regulatory role in mucosal immunity to actively reduce inflammatory responses (41, 115). We propose that increased TGF-β1 concurrent with high MUC2 mRNA expression and trafficking signaled by SEC31A, cTAGE5 and TANGO1 levels could reflect efforts to start reparative processes to restore the broken mucosal barrier (107, 111).

### Conclusion

This study is the first to describe that mucin-2 ER-to-Golgi transport occurs in large COPII vesicles, similar to the transport of other large proteins, such as PC1 and chylomicrons. The enlargement of the COPII vesicles for mucin-2 transport is facilitated by cTAGE5 and TANGO1 and promoted by Cullin-3 adaptor KLHL12. Successful generation of these large COPII vesicles is essential to maintain ER homeostasis and cellular function. IBD, an intestinal disease associated with loss of mucous layer and decreased TGF-β protein levels, presents increased ER stress levels of unknown cause. We demonstrated that TGF-β inhibition led to increased ER stress levels, as a result of decreased TANGO1 protein expression. Interestingly, TGF-β inhibition upregulated TANGO1 mRNA levels, which we hypothesize to be a mechanism of the UPR to ameliorate ER stress. Moreover, the discrepancy between TANGO1 mRNA and protein levels is suggestive of post-transcriptional regulation, possibly by TGF-β, divulging a promising avenue for future research. Mucosal gene expression data from intestinal tissue of patients with IBD substantiated high ER stress levels and showed high MUC2 and associated COPII transport gene expression, however, in association with upregulated TGF-β1 mRNA expression. In conclusion, we propose that the unsuccessful formation of large COPII vesicles, as a result of the observed TGF-β inhibition by Smad7, could be one of the sources of ER stress in IBD, because of lowered TANGO1 protein expression, subsequently leading to decreased mucin-2 secretion.

## Materials and Methods

### Cell Culture, Transfections, and Plasmids

HT29-MTX cells were cultured in Dulbecco’s Minimum Eagle Media F-12 (DMEM, Gibco, Cat no. 11320033) containing 2 mM GlutaMAX, supplemented with 10% Fetal Bovine Serum (FBS, Gibco, Cat no. 10270106), 100 units/mL of penicillin (Gibco, Cat no. 15140130), and 100 µg/mL of streptomycin (Gibco, Cat no. 15140130), and 1% non-essential amino acids (NEAA, Gibco, Cat no. 11140-050). Cells were maintained at 37 °C, 100% humidity, and 5% CO_2_ and were passaged twice per week using DBPS (Gibco, Cat no. 14190-094) and Trypsin-EDTA (Gibco, Cat no. 15400054). Starvation cell culture medium contained DMEM F-12 supplemented with 2 mM GlutaMAX, 100 units/mL of penicillin, 100 µg/mL of streptomycin, and 1% NEAA.

Cells were seeded at 15 000 cells/cm^2^ in a 6 wells plate (Greiner Bio-One, Cellstar, Cat no. 657 160) and cultured for 2 days. Four hours prior to transfection, the medium was changed to starvation medium. Subsequently, silencing RNAs (siRNA, Ambion Life Technologies, 25 pmol per well) were diluted in starvation medium. DharmaFect 1 (DharmaCon, 7.5 µL per well, Cat no. T-2001-02) was diluted in starvation medium as well and combined in a 1:1 ratio with the siRNA. The mixture was incubated for 20 minutes at room temperature before addition to the cells. Cells were transfected for approximately 40 hours after samples were collected for further analysis. The Silencer Pre-Designed siRNAs (Ambion) used were SEC31A (147540), TANGO1 (UNQ6077, 124540), cTAGE5 (144132), and KLHL12 (133405). A non-targeting siRNA (ASO2CMLU) was used as negative control for the silencing experiments.

The KLHL12-FLAG in pcDNA5 construct was a kind gift from Dr. Michael Rape (University of California, Berkeley, CA, USA) and the TANGO1-FLAG plasmid in pcDNA3.1 was kindly provided by Dr. Vivek Malhotra (CRG-Centre de Regulacio Genomica, Barcelona, Spain). E. Coli bacteria containing KLHL12-FLAG plasmid were cultured on a LB Agar plate (Fisher BioReagents, Cat no. BP1425-500) at 37 °C and the plasmid was isolated from single colonies using the PureYield Plasmid Midiprep kit (Promega, Cat no. A2495), according to the manufacturer’s instructions. Slbtl2 bacteria containing TANGO1-FLAG plasmid were handled similarly yet cultured at 30 °C. Plasmid concentrations were measured using the ND-1000 Spectrophotometer (NanoDrop, Unity Lab Services, Thermo Fisher Scientific) and the sequences were partially validated using Sanger Sequencing (Eurofins Genomics). Prior to transfection, 15 000 cells/cm^2^ were plated in a 6-wells plate and after one day transfected using Lipofectamine 2000 (Thermofisher, Cat no. 11668030) in OptiMEM (Gibco, Cat no. 31985-070) for 40 hours. The TANGO1-FLAG plasmid was transfected at 1 µg/mL, whereas the KLHL12-FLAG plasmid was transfected at 100 ng/mL, due to the formation of aggregates at higher concentrations. Transfection efficiency was determined by quantifying immunofluorescence using a FLAG antibody and DAPI counterstaining in ImageJ (v1.52p NIH).

It should be noted that the transfection efficiency of KLHL12-FLAG was lower than TANGO1-FLAG, due to using a ten times diluted plasmid and transfection agent (**Supplementary Figure 5**). However, the use of 1 µg of plasmid resulted in toxic aggregation of KLHL12, as observed by immunofluorescence of the FLAG protein (**Supplementary Figure 6A**). TANGO, a computational tool to predict beta aggregation of proteins, predicted beta aggregation for the KLHL12 protein (**Supplementary Figure 6B**) (116–118). Therefore, smaller amounts (0.1 µg) of KLHL12-FLAG were used to overexpress KLHL12 in HT29-MTX cells.

### Treatments

Brefeldin A (BFA) stock solution was prepared in DMSO at 500 µg/mL and Monensin stock solution was prepared in 70% ethanol at 1 mM. The cells were stimulated with BFA at 1 µg/mL, 10 µg/mL, or 100 µg/mL and Monensin at 1 µM for 45 minutes. Subsequently, the medium was refreshed to starvation medium for 24 hours after which samples were collected. Other treatments were incubated with the cells for 24 hours in starvation medium before sample collection. All treatments are summarized in **Table 1**.

**Table 1:**
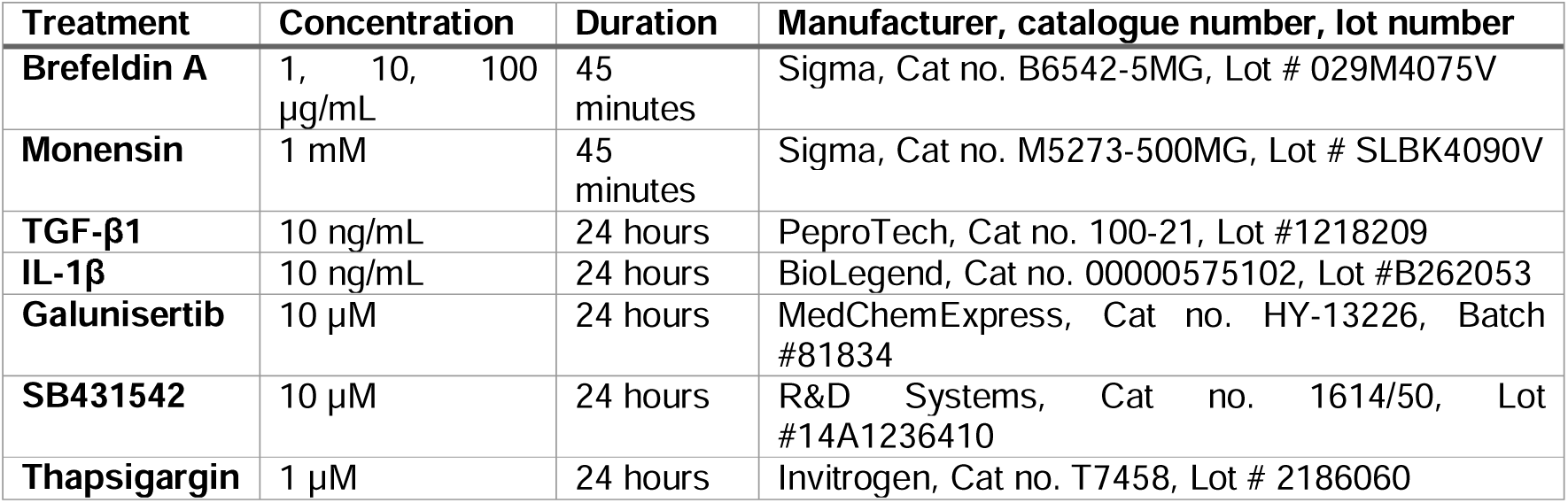
Treatments, concentrations, manufacturer, catalogue number, and lot number.

### Antibodies

Commercially available antibodies used for immunofluorescence (IF) and Western Blotting (WB) were as follows: mouse anti-mucin-2 (Santa Cruz, sc-7314, Lot #A3019, 1:200 for IF and 1:500 in TBST-XL for WB), rabbit anti-SEC31A (Cell Signaling Technologies, E1J5M, Lot #0, 1:800 for IF and 1:1000 in 5% Bovine Serum Albumin (Sigma, Cat no. A9418-50G) for WB), rabbit anti-TANGO1 (Sigma, Sigma, Cat no. HPA055922, Lot #000008249, 1:200 for IF, 1:1000 in 5% milk for WB), mouse anti-actin (Abcam, ab8224, 1:2000 in 5% milk for WB), mouse anti-FLAG (F1804-50UG, IF 1:400), rabbit anti-FLAG (Sigma, Cat no. F7425, Lot #06M4757, IF 1:100).

### Post-Golgi Mucin-2 Isolation for Western Blot

HT29-MTX cells were cultured in medium deprived of FBS 24h in 6-wells plates prior to mucus collection, as this avoided noisy Western Blot results and increased mucin-2 secretion, which was also observed by Cantero-Recasens et al. in HT29 cells (119), without decreasing cell viability (**Supplementary Figure 7**). After aspiration of medium, the cell layers were washed with 200 µL of 6 M Guadinium Hydrochloride (Gdn-HCl, Sigma, Cat no. G3272-25G), 5 mM EDTA (Fluka, Cat no. 03690-100ML), and 10 mM NaH_2_PO_4_ (Sigma, Cat no. 331988) at pH 6.5, and aspirated after 1 minute, loosely based on the method of Cantero-Recasens *et al*. (119). Subsequently, 1 mL of DBPS was added and incubated for 30 minutes at 37 °C, which was then carefully collected and precipitated using methanol/chloroform protein precipitation. Briefly, 4 mL of methanol (Sigma, Cat no. 322415-2L), 1 mL of chloroform (Sigma, Cat no. 288306), and 3 mL of ultrapure H_2_O (MilliQ, Salmenkipp, Purelab Flex, Elga) were added in that order and each to mix vortexed for 2 seconds. Subsequently, the samples were centrifuged for 5 minutes at 4000 rpm and the top aqueous layer was removed, as the protein was residing between the layers. Next, 4 mL of methanol was added, vortexed, and centrifuged for 14 minutes at 4000 rpm. The methanol was removed, and the protein pellet was dried using N_2_ air flow and dissolved in 12 µL MilliQ and 8 µL 5x Laemmli Buffer, consisting of 1.5 M Tris (Thermo Fisher, Cat no. 1.08382), 10% SDS (Sigma-Aldrich, L3771-500G), 0.05% Bromophenol Blue (Bio-Rad, Cat no. 1610-404), 50% glycerol (CalBiochem, Cat no. 356350-1000ML), and 1.43 M β-mercaptoethanol (Thermo Fisher, Cat no. 31350010). Samples were used directly for SDS-PAGE and Western Blot purposes or stored at -80 °C until further use.

### Cell Lysate Sample Preparation

Cell lysates were obtained from different serum-deprived 6 wells plates, as the Gdn-HCl mucin-2 extraction interfered with obtaining proper cell lysates. Samples were washed twice with ice cold DBPS and subsequently lysed in 200 µL RIPA buffer (50 mM Tris, 1% Triton X-100, 12 mM sodium deoxycholate (Sigma, Cat no. 30970-25G), 0.1% (w/v) SDS, 150 mM NaCl (Sigma, 746398-500G) in MilliQ), containing 1X Proteinase Inhibitor Cocktail (Calbiochem, Cat no. 539134). Subsequently, the cell lysate was collected and agitated for 30 min at 800 rpm at 4 °C in a Thermomixer (Eppendorf). Afterwards, the lysate was sonicated (42 kHz, Branson 1510, Bransonic) in an ice bath intermittently 3 times for 10 seconds each, maintaining 1 minute at 4 °C in between sonication pulses. Lastly, the sample was centrifuged for 20 minutes at 12000 rpm and 100 µL of the supernatant was used or stored at -80 °C until further use. Protein concentrations were measured using the DC Protein Assay (Bio-Rad, Cat no. 5000112) according to the manufacturer’s instructions.

### Western Blot

Mucin-2 precipitated samples could directly be processed, whereas 17.5 µg protein of cell lysates were combined with 8 µL of 5x Laemmli buffer. Protein samples were denatured at 100 °C for 5 minutes and briefly centrifuged before loading onto a 4-12% Tris-Glycine Novex Wedgewell gel (Fisher Scientific, Cat no. XP04125BOX). SDS-PAGE was performed using standard running buffer in Xcell Surelock Electrophoresis Cell (Invitrogen) at 150V for 75 minutes. Protein transfer was completed in a transfer buffer, containing 20% Emsure methanol, 0.1% (w/v) SDS, 25 mM TRIS-Base (Sigma, Cat no.), and 19.2 mM Glycine (Sigma, Cat no. G7126) in MilliQ for 100 minutes at 100 V onto a 0.45 µm PVDF membrane (Immobilon). Protein transfer was confirmed using a Coommassie Blue stain (Fisher Scientific, Cat no. 23236). The membrane was incubated in blocking buffer (5% non-fat dry milk (Carlroth, Cat no. T145.1) in TBST containing 0.005% Tween-20 (Sigma, Cat no. P7949-500ML), or 0.02% Tween-20 (TBST-XL)), for 1 hour at room temperature on an orbital shaker at 50 rpm. Primary antibodies were incubated overnight at 4 °C on an orbital shaker, as mentioned in section 2.3. The next morning, membranes were washed 3 times for 5 minutes in TBST or TBST-XL for mucin-2 and incubated with anti-mouse IgG HRP Conjugate (1:10000, Promega, Cat no. W4021) or anti-rabbit IgG HRP Conjugate (1:10000, Promega, Cat no. W4011) in blocking buffer and developed using the SuperSignal West Femto Maximum Sensitivity Substrate kit (Thermofisher Scientific, Cat no. 34095). Membranes were imaged using FluorChem M Imager (ProteinSimple) and analyzed in ImageJ (v1.52p, NIH).

### Immunofluorescence

Cells were seeded and cultured at 15 000 cells/cm2 in a 24 wells plate onto a #1.5 coverslip with a diameter of 12 mm (Thermo Scientific, Cat no. 1217N79) and deprived of serum for 24 hours before fixation. After 3 days, the cells were fixated using 4% formaldehyde (Sigma, Cat no. 252549) in DPBS for 15 minutes at room temperature and subsequently washed three times for 5 minutes with DPBS. Afterwards, samples were permeabilized in 0.1% (v/v) Triton X-100 (Sigma-Aldrich, Cat no. T8787-250ML) for 10 minutes at room temperature and blocked using the blocking buffer consisting of 5% (v/v) FBS in DBPS for 1 hour at room temperature under gentle shaking. The primary antibodies were diluted in blocking buffer and incubated overnight at 4 °C. The next day, samples were washed three times for 5 minutes in DPBS. The secondary antibodies (Goat anti-Mouse IgG Alexa Fluor 488, Invitrogen; Goat anti-Rabbit IgG Alexa Fluor 647, Invitrogen) were diluted 1:500 in blocking solution and incubated for 60 minutes at room temperature in the dark under gentle shaking. Subsequently, three washes of 5 minutes in DPBS were performed. The nuclei were stained using DAPI (1:4000 in DPBS, Invitrogen, Cat no. D1306) and the samples were mounted with DAKO mounting medium (Agilent, Cat no. S302380-2) onto a glass slide (Thermofisher, Cat no. 28908). Samples were imaged using the Fluorescent EVOS Microscope (Invitrogen EVOS FL Imaging System, ThermoFisher Scientific) or the Zeiss LSM 880 Confocal Microscope (Zeiss). F-actin immunostaining was performed by incubating fixated, permeabilized, and blocked cells with Phalloidin Alexa Fluor 633 (1:40 in DBPS, Invitrogen, Cat no. A22284) for 30 minutes at room temperature under gentle shaking. Subsequently, cells were washed 3 times with DPBS, counterstained with DAPI, and imaged using a fluorescent microscope.

### RT-qPCR

mRNA was isolated from (treated) cells in a 12 wells plate (Greiner Bio-One, Cellstar, Cat no. 665 180) using the NucleoSpin RNA kit (Macherey-Nagel, Cat no. 740955.250) according to the manufacturer’s instructions. RNA quality was ensured with 260/230 ratio between 1.8 and 2.2 using a ND-1000 spectrophotometer. cDNA samples were obtained from 1 µg RNA using the iScript^TM^ cDNA synthesis kit (Bio-Rad, Cat no. 170-8891) according to the manufacturer’s instructions. Reverse transcriptase quantitative polymerase chain reaction (RT-qPCR) was performed with 5 ng cDNA per sample, 10 µM of forward and reverse primer, and 2x SensiMix^TM^ SYBR & Fluorescein (Bioline, Cat no. QT615-05) in a 384 wells plate (Bio-Rad, Cat no. HSP3805). Thermal cycling was performed in a Bio-Rad CFX384 thermocycler using 40 cycles of 95 °C melting for 15 seconds, 60 °C annealing for 15 seconds, and 72 °C extension for 15 seconds, prior to an initial hot start at 95 °C for 10 minutes (see **Supplementary Table 1** for primer sequences).

### Cell Viability

Cell viability was measured by measuring ATP production using Cell Titer Glo assay 2.0 (Promega, Cat no. G9241). Cells were seeded at a density of 15 000 cells/cm^2^ in an opaque 96 well plate (Greiner Bio-one, Cat no. 655098) in 100 µL cell culture medium. After 3 days of culture and relevant cell treatments, a volume of 100 µL of reagent was added to the cell culture medium and allowed to react for 10 minutes at room temperature, under slow orbital shaking (50 rpm, KS250 Basic, Ika Labortechnik). Staurosporin (10 µM, Millipore Sigma, Cat no. 569397-100UG) was used as a negative control. Luminescence was measured using Victor3 1420 Multilabel Counter (PerkinElmer) for 1 second per sample.

### Clinical validation in a cohort of individuals with inflammatory bowel disease (IBD)

A human validation of the *in vitro* results was performed using intestinal biopsy samples from individuals diagnosed with inflammatory bowel disease (IBD). Participants were recruited from the outpatient clinic of the University Medical Center Groningen (UMCG) as part of the 1000IBD project, which involved gathering comprehensive phenotypic information and molecular profiles (120). Patients that were included were >18 years of age, enrolled in the period of 2003 until 2019, and were diagnosed with Crohn’s disease (CD) or ulcerative colitis (UC). The diagnosis of IBD was based on various criteria, including clinical symptoms, laboratory tests, endoscopic examination, and histopathological analysis of biopsies. Additionally, biopsy samples from 16 healthy individuals without IBD were included for comparison. Their biopsies were obtained during endoscopies conducted due to suspected intestinal disease or as part of colon cancer screening, all of which yielded negative results. Prior to sample collection, written informed consent was obtained from all participants. The study was approved by the Institutional Review Board (IRB) of the UMCG in Groningen, the Netherlands (in Dutch: ’Medisch Ethische Toetsingscommissie’, METc; IRB nos. 2008/338 and 2016/424), and adhered to the principles outlined in the Declaration of Helsinki (2013).

### Collection of intestinal biopsies

A total of 711 intestinal biopsies were obtained from 420 patients diagnosed with IBD along with 52 intestinal biopsies from 16 healthy non-IBD controls (121). During an endoscopic procedure, biopsies were immediately frozen in liquid nitrogen by either an endoscopy nurse or a research technician. Biopsies were taken from both inflamed and non-inflamed regions in adjacent areas, and their inflammatory status was assessed histologically by certified pathologists. For patients with CD and UC, biopsies were taken from both the ileal and colonic tissue. However, inflamed ileal biopsies from UC patients (n=3) were excluded from the analysis, likely due to backwash ileitis. The biopsies were stored at -80°C until further processing.

### RNA isolation, sequencing, and data processing

RNA isolation was conducted using the AllPrep DNA/RNA mini kit (Qiagen, ref no: 80204) following the manufacturer’s instructions. To homogenize the intestinal biopsies, the Qiagen Tissue Lyser with stainless steel beads (diameter 5 mm, reference number: 69989) was utilized in RLT lysis buffer containing β-mercaptoethanol. In the first batch of samples, the BioScientific NEXTflexTM Rapid Directional RNA-Seq Kit (Perkin-Elmer) was employed for sample preparation. Paired-end RNA sequencing was performed on the Illumina NextSeq500 sequencer (Illumina). For the second batch of samples, the Eukaryotic Transcriptome Library (Novogene) was constructed for sample preparation, followed by paired-end RNA sequencing on the Illumina HiSeq PE250 platform. Sequencing was carried out in two separate batches, and pseudo-randomization was employed across plates to mitigate potential batch effects, taking into account the type of IBD diagnosis, biopsy location, and disease activity. Batch effects were considered in all the analyses. On average, approximately 25 million reads were generated per sample. The quality of raw reads was assessed using FastQC with default parameters (ref v.0.11.7). Adapters identified by FastQC were removed using Cutadapt (ref v1.1) with default settings. Sickle (ref v1.200) was used to trim low-quality ends from the reads (length <25 nucleotides, quality <20). Alignment of the reads to the human genome (human_glk_v37) was performed using HISAT (ref v0.1.6) with a maximum allowance of two mismatches, and read sorting was done using SAMtools (ref v0.1.19). SAMtools flagstat and Picard tools (ref v2.9.0) were utilized to obtain mapping statistics. Six samples with a low percentage of read alignment (<90%) were excluded from the analysis. Gene expression was estimated using HTSeq (ref v0.9.1) based on Ensemble version 75 annotation, resulting in an RNA expression dataset comprising 15,934 genes. Gene level expression data were normalized using a trimmed mean of M values, and clr transformation was applied, resulting in a set of 826 mucosal RNA-seq samples.

### Statistical Analysis

Evaluation of statistical significance was determined using a one- or two-way ANOVA followed by Bonferroni’s post-hoc test using GraphPad Prism 8.0.1 (GraphPad Software, La Jolla, CA). A p-value <0.05 was considered to indicate statistical significance. All experiments were conducted in triplicates (n = 3) unless otherwise stated. For human intestinal mucosal RNA-sequencing analysis, gene expressions were compared between group of individuals with CD, UC and controls using Mann-Whitney *U*-tests followed by post-hoc Bonferroni correction for multiple testing. Analyses and visualization of gene expressions were performed using the Python programming language (v.3.9.0; Python Software Foundation; https://python.org) using the *pandas* (v.1.4.2), *numpy* (v.1.21.0), *matplotlib* (v.3.5.1), and *seaborn* (v.0.12.1) packages.

## Acknowledgments

We thank Dr. Michael Rapé (University of California, Berkeley, CA, USA) for kindly gifting the KLHL12-FLAG in pcDNA5 construct and Dr. Vivek Malhotra (CRG-Centre de Regulacio Genomica, Barcelona, Spain) for kindly providing the TANGO1-FLAG plasmid in pcDNA3.1.

## Author Contributions

Margaretha A.J. Morsink – Conceptualization; Data curation; Formal analysis; Validation; Investigation; Visualization; Methodology; Carried out the research and collected and analyzed the data; Writing—original draft;

Lena S. Koch – Conceptualization; Resources; Data curation; Validation; Visualization; Investigation; Formal analysis; Methodology; Supervision; Writing – review and editing; Writing – final draft

Shixian Hu – Clinical validation: Conceptualization; Data curation; Formal analysis; Validation; Investigation; Visualization; Methodology;

Rinse K. Weersma – Clinical validation: Conceptualization; Resources; Supervision; Funding acquisition; Data curation; Project administration;

Harry van Goor – Clinical validation: Clinical validation: Conceptualization; Data curation; Formal analysis; Validation; Investigation; Visualization; Methodology;

Arno R. Bourgonje – Clinical validation: Clinical validation: Conceptualization; Data curation; Formal analysis; Validation; Investigation; Visualization; Methodology;

Kerensa Broersen - Conceptualization; Resources; Supervision; Funding acquisition; Data curation; Project administration; Writing – original draft; Writing—review and editing

## Supplementary Figures

**Supplementary Figure 1:**
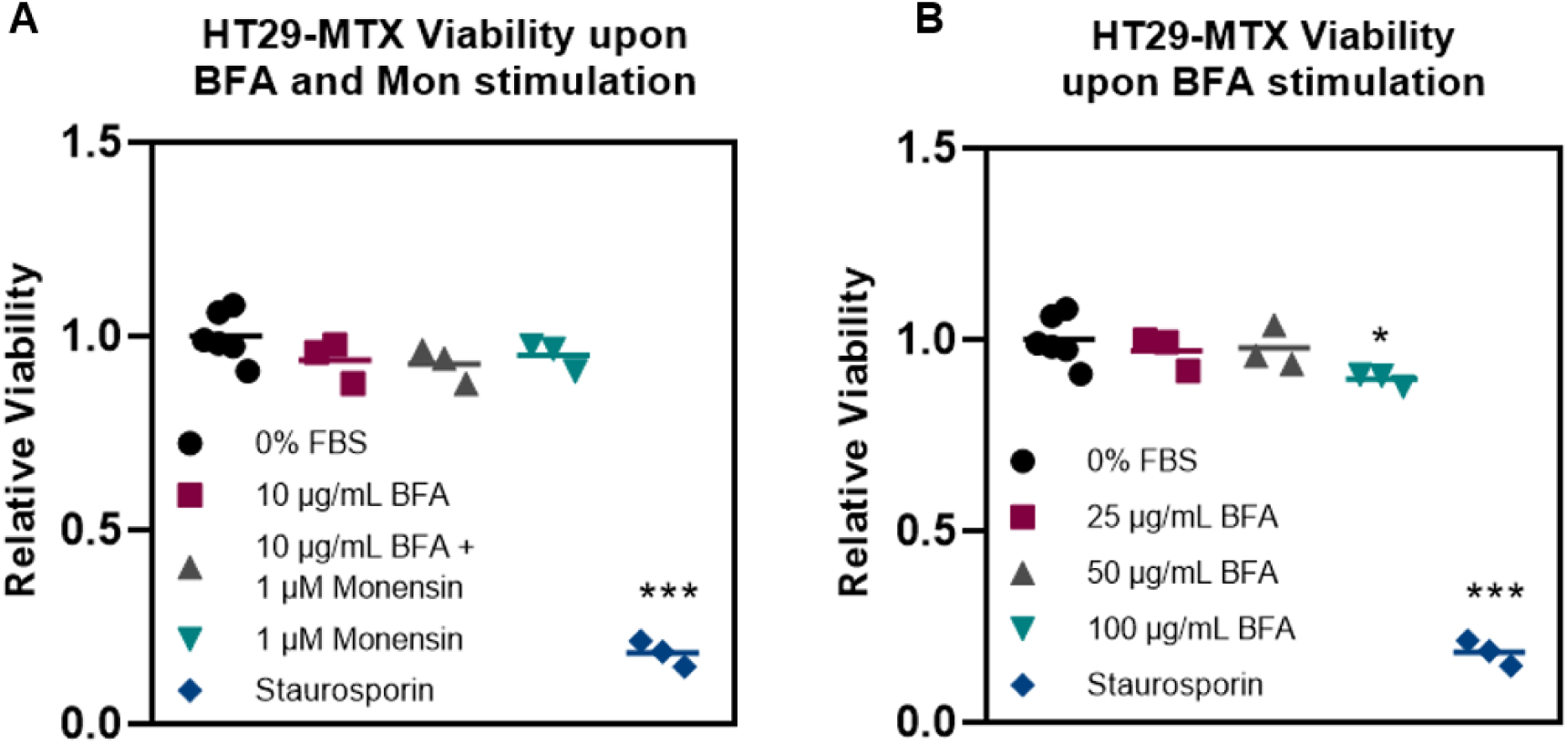
ATP production as measure of viability of HT29-MTX cells upon addition of BFA and Monensin. A) Concentrations of BFA and Monensin used to determine conventional secretion of MUC2 (10 µg/mL of BFA and 1 µM of Monensin). B) Concentrations of BFA used to determine BFA-dose dependency (25 µg/mL; 50 µg/mL; 100 µg/mL). Staurosporin (10 µM) was used as a negative control. Data was analyzed using one-way ANOVA followed by Bonferroni’s test. Significance was indicated as * (p < 0.05); ** (p < 0.01); *** (p <0.001) with respect to 0% FBS.

**Supplementary Figure 2:**
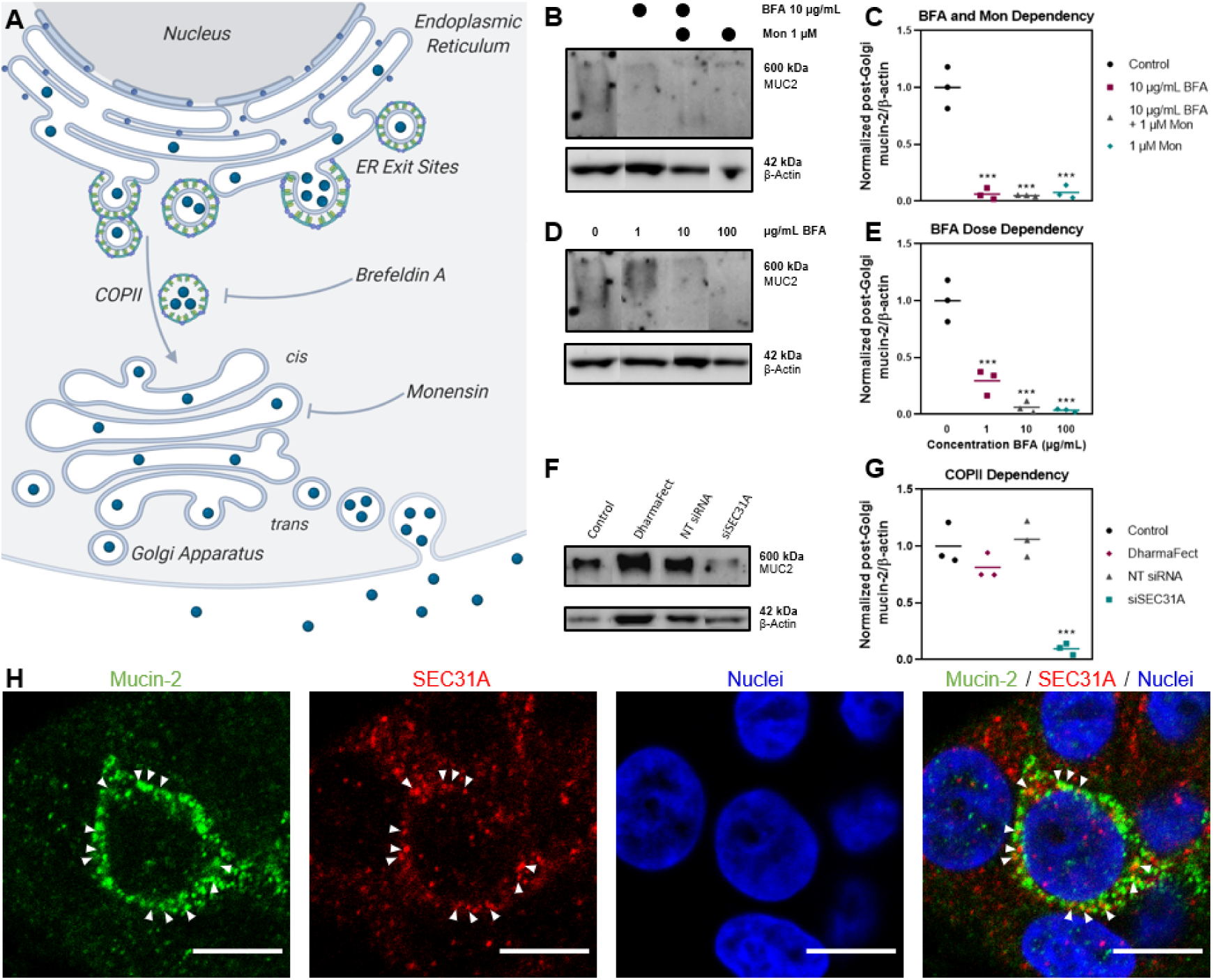
Mucin-2 is trafficked in a COPII-dependent manner. A) A schematic representation of the effect of Brefeldin A (BFA) and Monensin (Mon) on conventional protein secretion. Created with BioRender.com. B) Western Blot of post-Golgi mucin-2 as a result of BFA and Mon exposure for 45 minutes. C) Semi-quantification of the Western Blot using ImageJ, comparing the signal intensity of mucin-2 to the signal intensity of β-actin, indicating a statistically significant decrease of post-Golgi mucin-2 upon addition of BFA and Mon. D) Western Blot showing BFA dose dependency on post-Golgi mucin-2 levels. E) Semi-quantification of the Western Blot using ImageJ, comparing the signal intensity of mucin-2 to the signal intensity of β-actin, indicating a statistically significant decrease of post-Golgi mucin-2 secretion upon addition of different BFA concentrations. F) Western blot of silencing of COPII protein SEC31A showing decreased post-Golgi mucin-2 levels compared to controls, including non-targeting siRNA and transfection agent DharmaFect. G) Semi-quantification of the Western Blot using ImageJ, comparing the signal intensity of mucin-2 to the signal intensity of β-actin, indicating a statistically significant decrease of post-Golgi mucin-2 levels upon silencing of outer COPII protein SEC31A. H) Immunofluorescence of mucin-2 (green), SEC31A (red), and nuclei (blue), showing a partial colocalization of mucin-2 and SEC31A. Scale bar represents 10 µm. Data was analyzed using one-way ANOVA followed by Bonferroni’s test. Significance was indicated as * (p < 0.05); ** (p < 0.01); *** (p <0.001) with respect to Control (C and G) or 0 µM BFA (E).

**Supplementary Figure 3:**
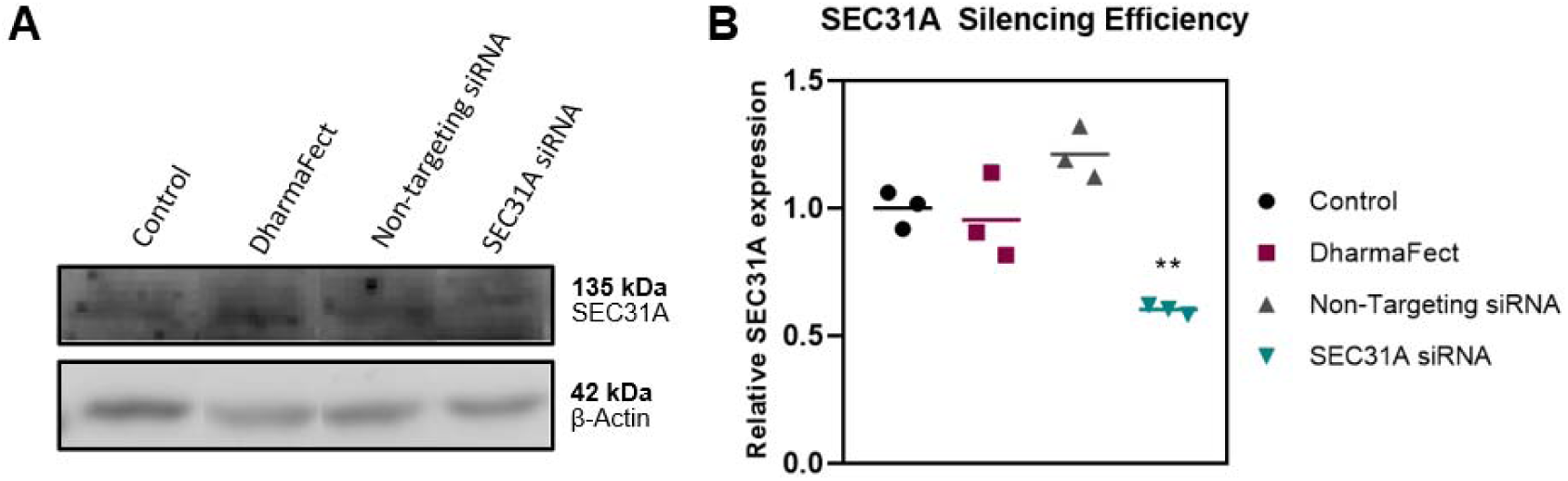
Silencing efficiency of SEC31A was determined at 39.72 ± 0.01882%. A) Western Blot data of SEC31A and β-actin as loading control for the control cells, the effect of DharmaFect, the effect of non-targeting siRNA, and the effect of SEC31A siRNA. B) Quantification of SEC31A expression using ImageJ, normalized to β-actin and control. Data was analyzed using one-way ANOVA followed by Bonferroni’s test. Significance was indicated as * (p < 0.05); ** (p < 0.01); *** (p <0.001) with respect to Control.

**Supplementary Figure 4:**
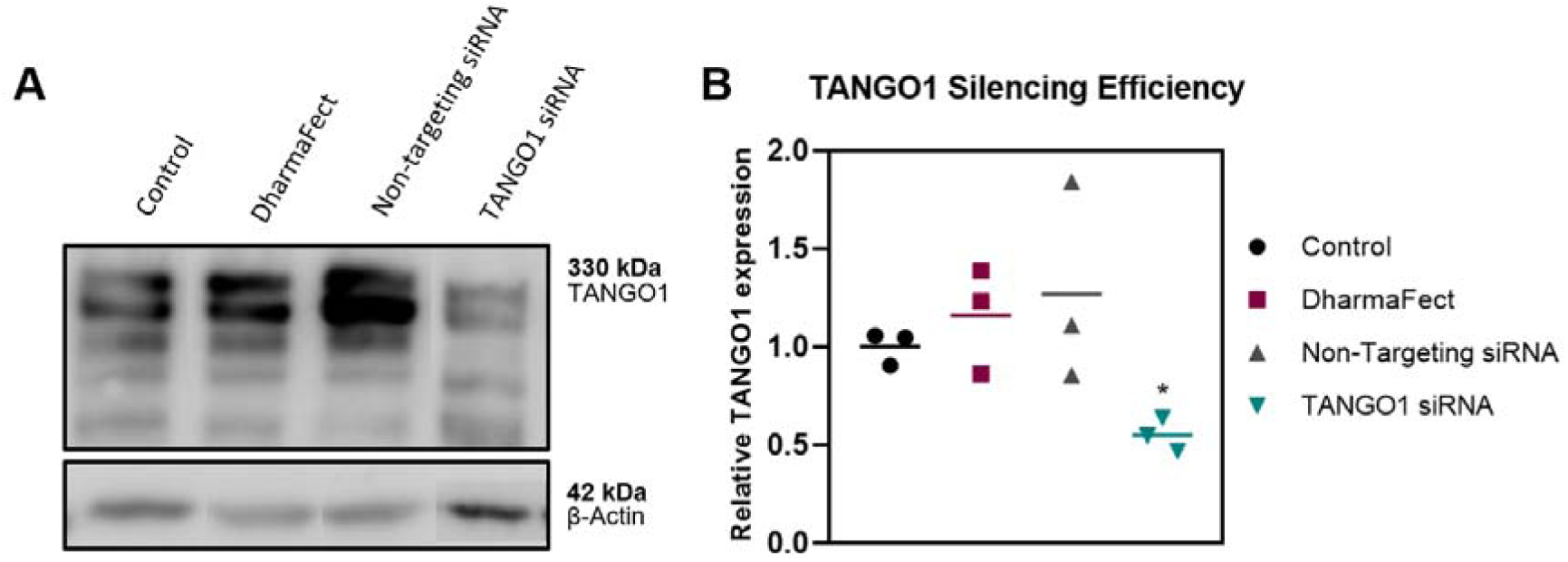
Silencing efficiency of TANGO1 was determined at 44.84 ± 0.085%. A) Western Blot data of TANGO1 and β-actin as loading control for the control cells, the effect of DharmaFect, the effect of non-targeting siRNA, and the effect of TANGO1 siRNA. B) Quantification of TANGO1 expression using ImageJ, normalized to β-actin and control. Data was analyzed using one-way ANOVA followed by Bonferroni’s test. Significance was indicated as * (p < 0.05); ** (p < 0.01); *** (p <0.001) with respect to Control.

**Supplementary Figure 5:**
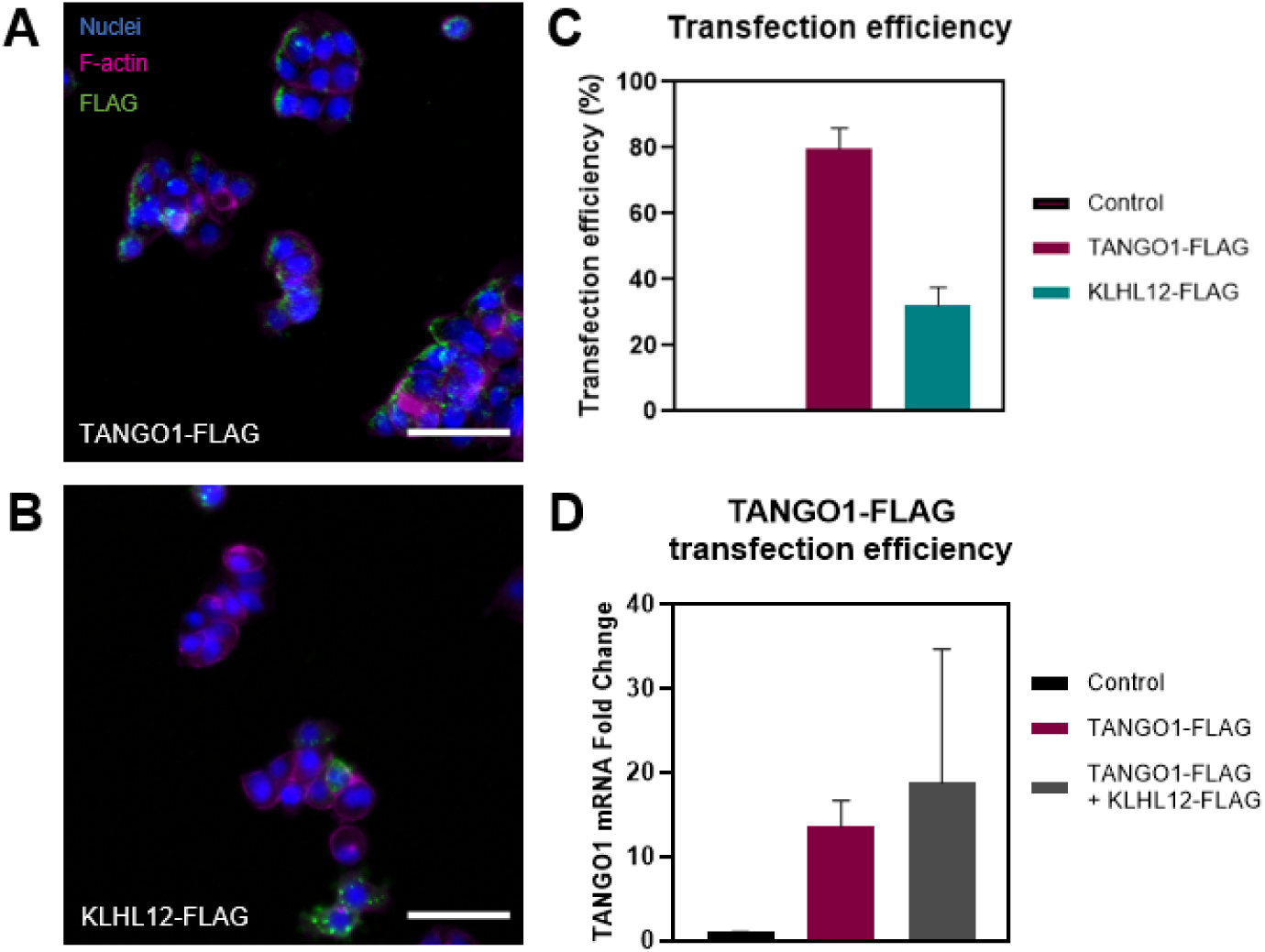
Transfection efficiency of TANGO1-FLAG and KLHL12-FLAG. A-B) Immunofluorescence of F-actin (Magenta), and nuclei (blue) and A) TANGO1-FLAG (green) and B) KLHL12-FLAG (green). Scale bar represents 50 µm. C) Transfection efficiency (%) of 80% for TANGO1-FLAG and 35% for KLHL12-FLAG. D) TANGO1 mRNA levels in transfected conditions.

**Supplementary Figure 6:**
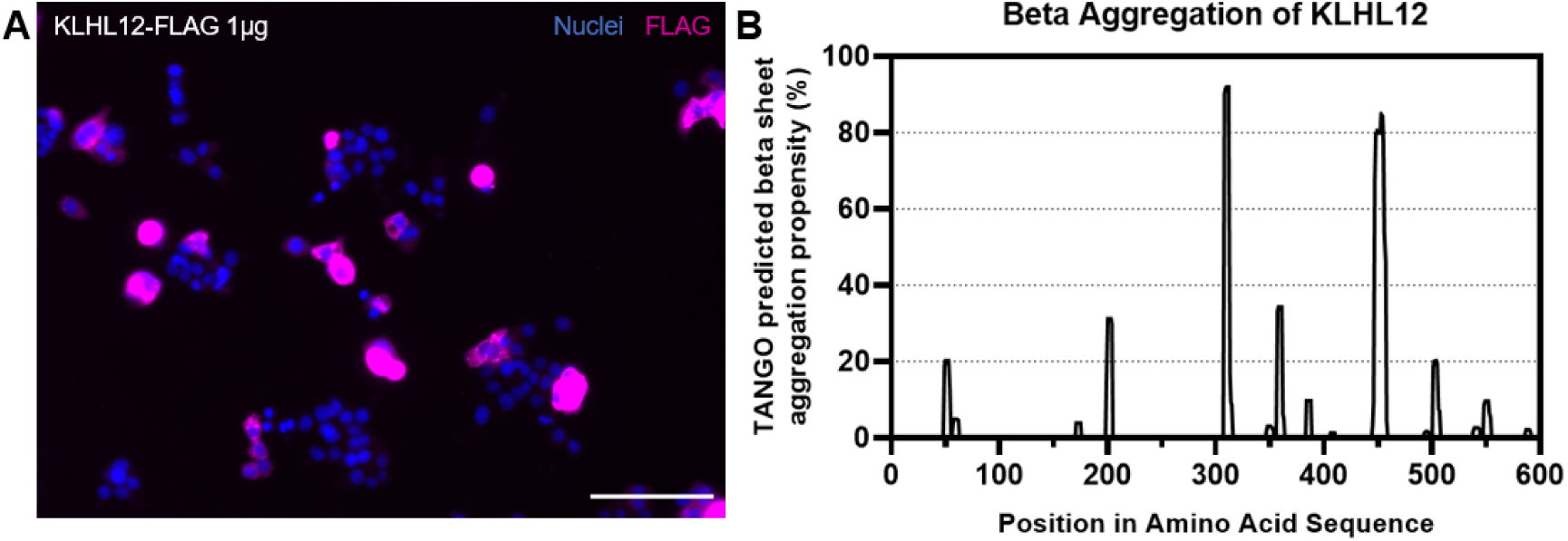
Toxic aggregation of KLHL12-FLAG by transfecting 1 µg. A) Immunofluorescent imaging of KLHL12-FLAG (Magenta) and nuclei (blue), showing large aggregates. Scale bar 100 µm. B) TANGO predicted beta sheet aggregation propensity indicating the aggregation of KLHL12 (116–118).

**Supplementary Figure 7:**
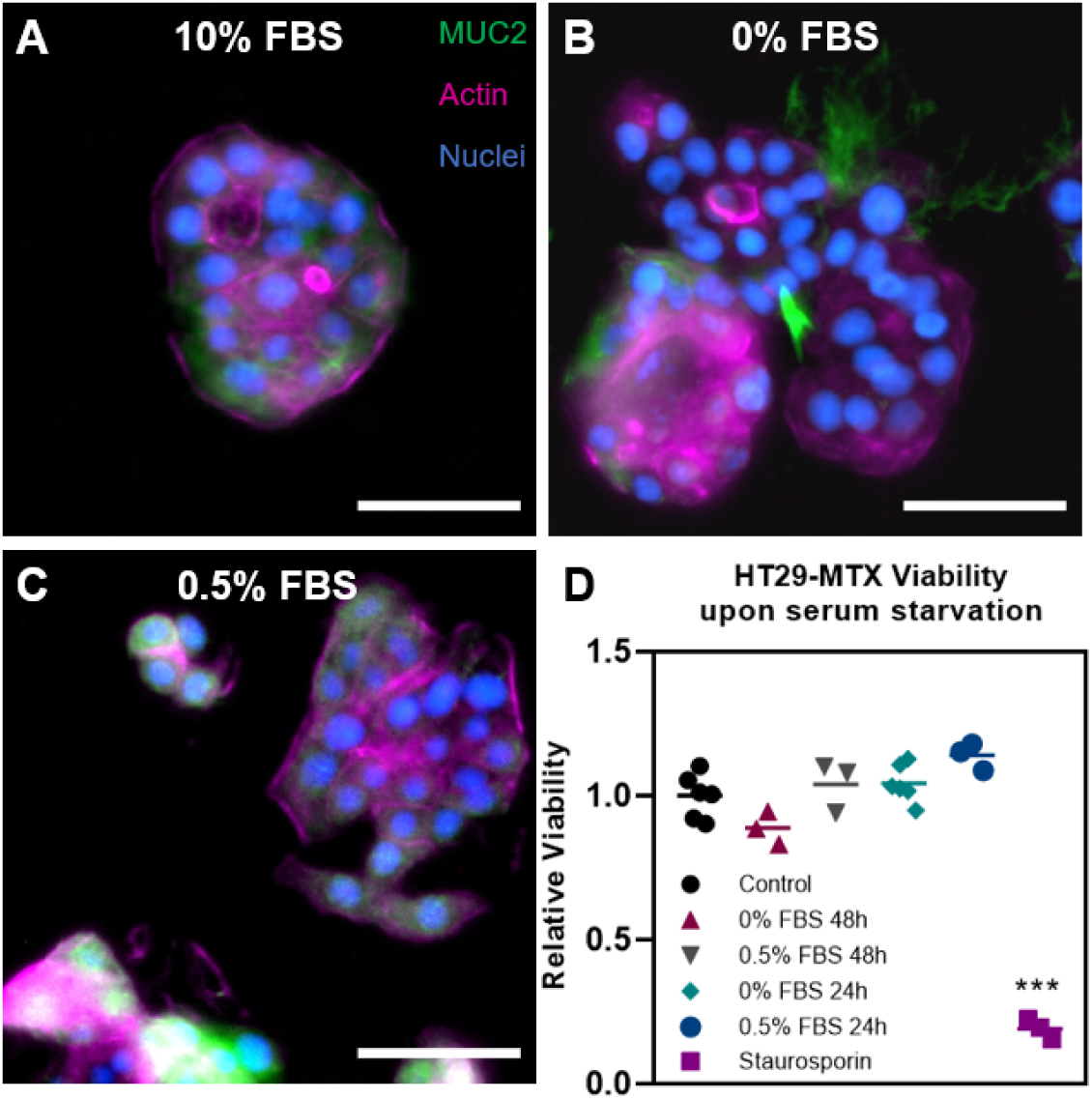
Response of HT29-MTX to different serum conditions. A-C) Immunofluorescence staining of HT29-MTX cells with MUC2 (green), F-actin (magenta), and nuclei (blue) for 10%, 0%, and 0.5% FBS for 24 hours respectively. Scale bar 50 µm. D) ATP production of HT29-MTX upon 10% FBS as positive control, and 0% and 0.5% FBS for 24h and 48h. Staurosporin (10µM) is used as negative control. Data was analyzed using one-way ANOVA followed by Bonferroni’s test. Significance was indicated as * (p < 0.05); ** (p < 0.01); *** (p <0.001) with respect to Control.

**Supplementary Figure 8:**
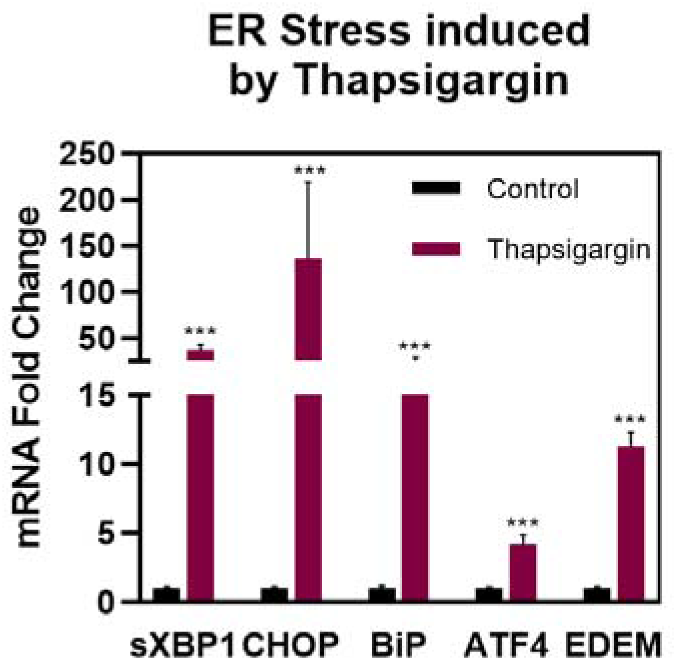
ER stress is measured by quantifying mRNA levels of sXPB1, CHOP, BiP, ATF4, and EDEM, as shown by stimulation with ER stressor Thapsigargin. Data was analyzed using one-way ANOVA followed by Bonferroni’s test. Significance was indicated as * (p < 0.05); ** (p < 0.01); *** (p <0.001) with respect to Control.

**Supplementary Table 1:**
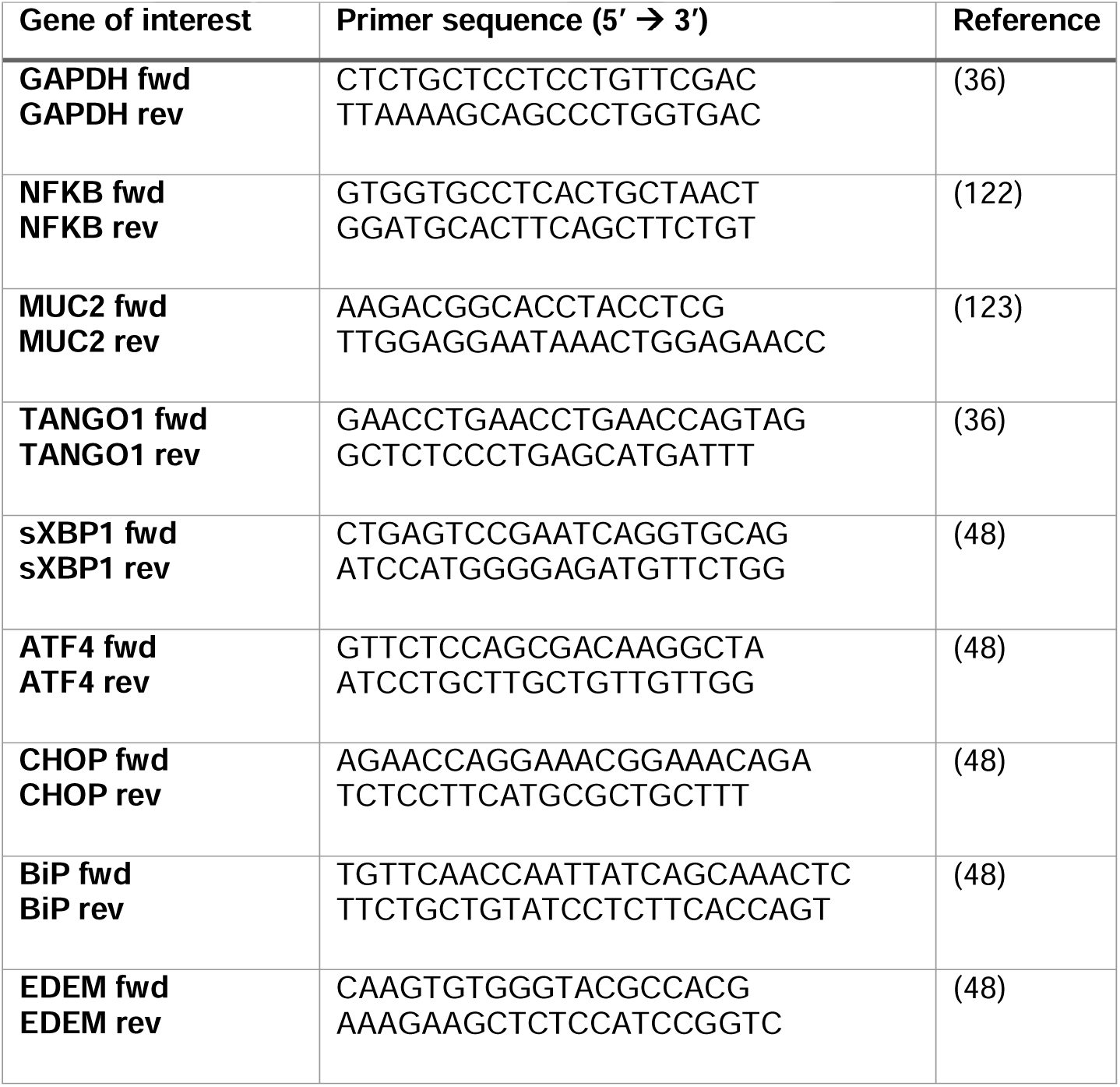
Primer sets for RT-qPCR.

